# PTP1B supports survival and regulates CFL1 and HSP90α abundance in trastuzumab-resistant HER2-positive breast cancer

**DOI:** 10.64898/2026.10.23.761095

**Authors:** Vinay Mandati, Maria C. Panepinto, Christy Felice, Sean Krivitsky, Suzanne Russo, Payal Naik, Raditya Utama, David L. Spector, Paolo Cifani, Nicholas K. Tonks

## Abstract

Trastuzumab resistance limits the treatment of HER2-positive breast cancer. Here, we identify protein tyrosine phosphatase 1B (PTP1B) as a survival dependency in models of intrinsic and acquired trastuzumab resistance, including cells with endogenous *ERBB2* kinase domain mutations (L755S and V777L), a patient-derived tumor organoid model, and orthotopic xenografts. PTP1B expression was elevated in multiple trastuzumab-resistant models, accompanied by increased PI3K–AKT and ERK signaling in several of these. The PTP1B inhibitor DPM-1003 suppressed growth of trastuzumab-resistant cells and organoids, inhibited xenograft progression, and induced apoptosis. Genetic ablation of PTP1B reduced growth and restored trastuzumab responsiveness in multiple resistance models, although JIMT1 cells remained resistant. Quantitative phosphoproteomics, phosphotyrosine immunoprecipitation, and substrate-trapping assays identified cofilin-1 (CFL1) and HSP90α as substrates of PTP1B. Loss of PTP1B increased their tyrosine phosphorylation, reduced their abundance, and increased their recovery in ubiquitin pull-downs. Proteasome inhibition restored substrate abundance, and PTP1B re-expression restored protein levels in PTP1B-null cells. These findings support a model in which PTP1B-dependent dephosphorylation limits turnover of the substrates CFL1 and HSP90α. Genetic or pharmacological targeting of PTP1B suppressed survival-associated signaling and increased apoptotic markers. Together, these results identify PTP1B as a therapeutic vulnerability in trastuzumab-resistant HER2-positive breast cancer and implicate regulation of CFL1 and HSP90α abundance in the accompanying signaling response.

**Significance:** PTP1B targeting suppresses growth of trastuzumab-resistant HER2-positive breast cancer models and implicates phosphatase-dependent control of substrate turnover as a potential mechanism supporting tumor cell survival and therapeutic resistance.

## Introduction

Targeted therapies have transformed cancer treatment, but therapeutic resistance often limits their efficacy (1–6). In HER2-positive breast cancer, trastuzumab has significantly improved patient outcomes, yet many tumors exhibit intrinsic resistance or relapse following an initial response (6–13). HER2-driven tumors depend on PI3K–AKT and RAS–MAPK signaling for proliferation and survival, with HIF-1α-associated responses also contributing to tumor cell fitness (14–16). Diverse resistance mechanisms preserve these signaling networks during treatment, highlighting the need to identify shared regulators of the resistant state (13,17).

Trastuzumab resistance involves both pre-existing alterations and adaptive signaling changes. Resistance-associated alterations include *PIK3CA* mutations (18), *PTEN* loss (19), truncated p95HER2 (20), and activating *ERBB2* kinase-domain mutations such as L755S and V777L, which can alter sensitivity to HER2-targeted agents (21–25). Acquired resistance can also involve compensatory signaling through EGFR, HER3, MET, and IGF-1R (26–30). These mechanisms can converge on sustained proliferation and survival, creating potential vulnerabilities beyond HER2 itself.

Reversible tyrosine phosphorylation is controlled by protein tyrosine kinases and phosphatases (PTPs). Although anticancer drug development has largely focused on kinases, PTPs can also promote oncogenic signaling and represent therapeutic targets (31–38). Their biological effects depend on the substrates and signaling networks engaged in a particular cellular context. Among these, Protein Tyrosine Phosphatase 1B (PTP1B) has emerged as an important regulator of HER2-driven breast cancer. PTP1B is overexpressed in a subset of breast cancers and is associated with HER2 overexpression in human tumors (39). Genetic ablation or pharmacological inhibition of PTP1B delays HER2/Neu-driven mammary tumorigenesis and suppresses metastatic progression in preclinical models (40, 41). PTP1B promotes signaling through selective substrate dephosphorylation, including regulation of SRC family kinases, p62Dok-dependent RAS–MAPK signaling, and CDK3-dependent cell-cycle progression (42–48). Phosphoproteomic studies have also linked PTP1B to signaling proteins associated with breast cancer invasion and survival, including BCAR3, BCAR1, and CDCP1 (48–53). Furthermore, in HER2-positive cells, PTP1B regulates RNF213-dependent non-mitochondrial oxygen consumption and survival during hypoxic stress (54, 55). Whether these oncogenic and stress-adaptive functions contribute to trastuzumab resistance remains unresolved.

Here, we investigated PTP1B in intrinsic– and acquired-resistance cell lines, isogenic cells harboring endogenous *ERBB2* L755S or V777L mutations, a patient-derived tumor organoid model, and orthotopic xenografts. PTP1B expression was elevated in multiple resistance models, accompanied by increased AKT and ERK phosphorylation. Pharmacological inhibition suppressed resistant cell and tumor growth and induced apoptosis, whereas genetic deletion restored trastuzumab responsiveness in selected models. Quantitative phosphotyrosine proteomics and substrate-trapping assays identified cofilin-1 (CFL1) and HSP90α as PTP1B substrates. PTP1B loss increased their tyrosine phosphorylation, reduced their abundance, and enhanced their recovery in ubiquitin pull-downs; proteasome inhibition restored protein levels. These findings support a model in which PTP1B-dependent dephosphorylation limits substrate degradation. PTP1B targeting also reduced survival-associated signaling, HIF-1α, BCL2, and Cyclin D1 expression and increased apoptotic markers. Together, these results identify PTP1B as a therapeutic vulnerability in trastuzumab-resistant HER2-positive breast cancer and implicate regulation of CFL1 and HSP90α stability in the accompanying signaling response.

## Results

### PTP1B expression was increased in models of intrinsic and acquired trastuzumab resistance

We compared intrinsically trastuzumab-resistant (HR) HER2-positive breast cancer cell lines JIMT1, HCC1419, HCC1569, and HCC1954 with sensitive (HS) BT474 and SKBR3 cells. Trastuzumab inhibited growth of the sensitive cells but had little effect in the resistant lines (**Figure 1A**), consistent with its predominantly cytostatic activity (56–59). Most intrinsic resistance models expressed more PTP1B than the sensitive controls, together with increased HIF-1α, phospho-AKT, and cyclin-D1 (**Figure 1B**). Analysis of DepMap datasets further supported these observations, demonstrating higher *PTPN1* expression (the gene encoding PTP1B) in trastuzumab-resistant than in trastuzumab-sensitive HER2-positive breast cancer cell lines (**Figure 1C**).

**Figure 1.**
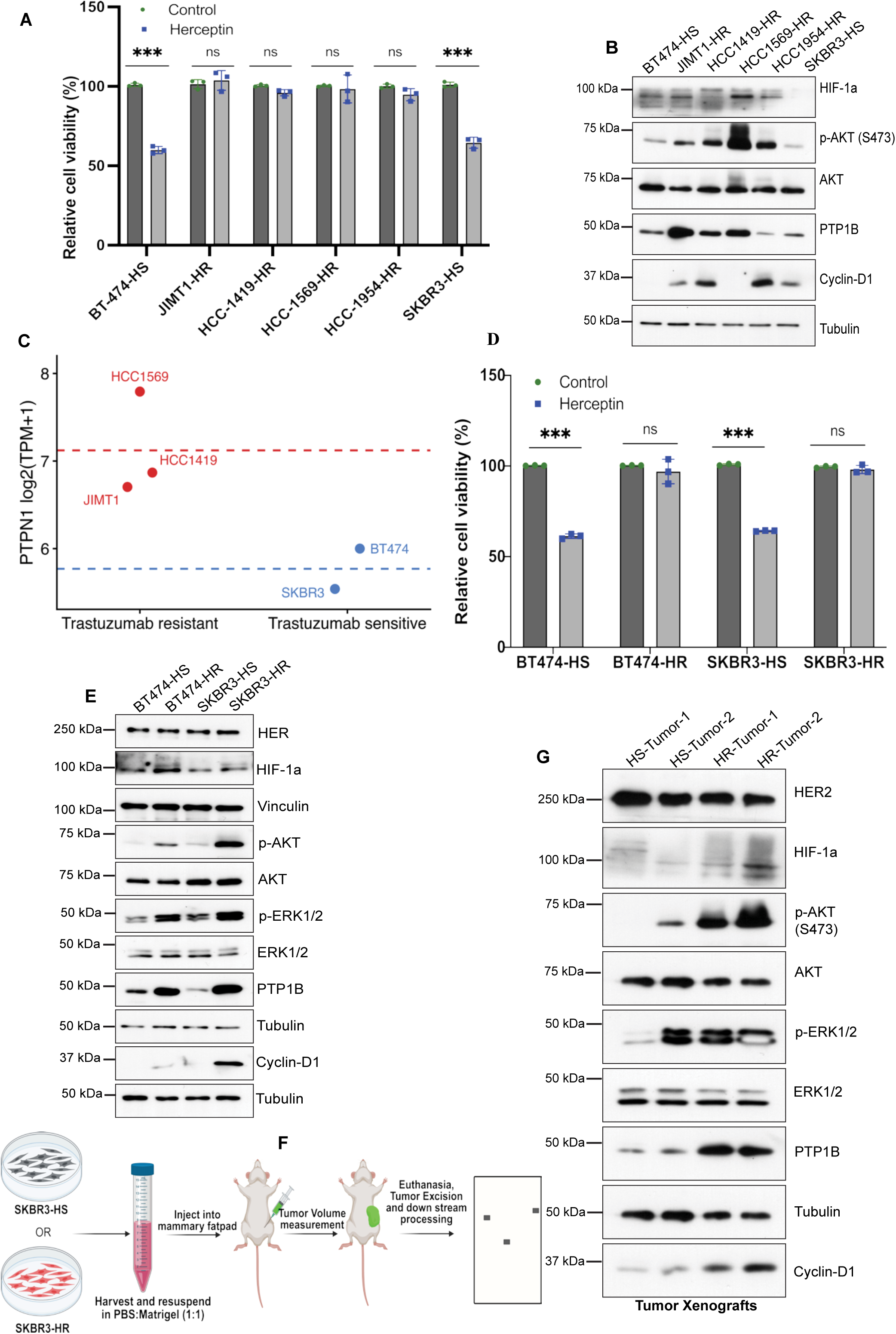
PTP1B was elevated in intrinsic and acquired models of trastuzumab resistance and was associated with sustained oncogenic signaling. **(A)** Cell viability measured by the CellTiter-Glo (CTG) assay in trastuzumab-sensitive (HS; BT474 and SKBR3) and intrinsic trastuzumab-resistant (HR; JIMT1, HCC1419, HCC1569, and HCC1954) HER2-positive breast cancer cell lines following trastuzumab treatment (15 μg/mL, 72 hrs). HS cells exhibited growth inhibition, whereas HR cells were largely resistant to treatment. **(B)** Immunoblot analysis of PTP1B, HIF-1α, and indicated signaling proteins in HS (BT474 and SKBR3) and intrinsic HR (JIMT1, HCC1419, HCC1569, and HCC1954) HER2-positive cell lines, showing elevated PTP1B expression, increased HIF-1α levels, sustained p-AKT and p-ERK signaling, and increased Cyclin D1 expression in resistance cells. **(C)** DepMap analysis of *PTPN1* transcript expression in HER2-positive breast cancer cell lines, presented as log₂(TPM + 1), where TPM denotes transcripts per million. Blue and red symbols represent cell lines categorized into lower– and higher-PTPN1-expressing groups, respectively. **(D)** CellTiter-Glo (CTG) assay of parental trastuzumab-sensitive (BT474-HS and SKBR3-HS) and acquired trastuzumab-resistant (BT474-HR and SKBR3-HR) cell lines following trastuzumab treatment (72 h), confirming acquisition of resistance. **(E)** Immunoblot analysis of PTP1B, HIF-1α, and indicated signaling proteins in HS and acquired HR BT474 and SKBR3 cells, demonstrating increased PTP1B expression, elevated HIF-1α levels, sustained p-AKT and p-ERK signaling, and increased Cyclin D1 expression in resistance cells. **(F)** Experimental scheme illustrating orthotopic implantation of SKBR3-HS and SKBR3-HR cells for xenograft generation and subsequent analysis of PTP1B and associated signaling proteins shown in **(G).** Created in BioRender. M, V. (2026) https://BioRender.com/sw1qf8n. **(G)** Immunoblot analysis of xenograft tumors derived from SKBR3-HS and SKBR3-HR cells showing increased PTP1B expression, elevated HIF-1α levels, sustained p-AKT and p-ERK signaling, and increased Cyclin D1 expression in resistance tumors. Data are presented as mean ± SD from three independent biological experiments unless otherwise indicated. Data were normalized to the respective control for visualization. Statistical significance in A and D was determined using two-way ANOVA followed by Bonferroni’s multiple comparisons test. ns, not significant; *P < 0.05; **P < 0.01; ***P < 0.001.

To determine whether PTP1B upregulation is also associated with acquired resistance, we generated trastuzumab-resistant derivatives of BT474 and SKBR3 cells through prolonged trastuzumab exposure using established protocols (50). The resulting BT474-HR and SKBR3-HR cells continued to proliferate in the presence of trastuzumab, confirming acquisition of resistance (**Figure 1D; Supplementary Figures S1A-S1D**). Similar to the intrinsic resistance models, acquired-resistance cells showed increased PTP1B expression and elevated pAKT and pERK, along with increased Cyclin D1 and HIF-1α expression, relative to their parental cells (**Figure 1E**).

Orthotopic SKBR3-HR xenografts also showed increased PTP1B and HIF-1α, together with higher AKT and ERK phosphorylation, compared with SKBR3-HS tumors (**Figures 1F–1G**). Cell viability assays further demonstrated increased growth-associated phenotypes in resistant cell models (**Supplementary Figures S1E–S1F**). These findings associate elevated PTP1B with trastuzumab resistance in both intrinsic and acquired models.

### ERBB2-mutant cells and trastuzumab-resistant tumor organoids exhibited increased PTP1B expression

ERBB2 kinase-domain mutations can alter oncogenic signaling and responses to HER2-targeted agents (21, 60–62). We examined L755S and V777L (**Figure 2A**), which have been associated with distinct responses to trastuzumab and lapatinib; L775S, one of the more frequently observed kinase-domain mutations, favors the active conformation of HER2 and has been associated with lapatinib resistance (63), whereas V777L enhances HER2 kinase activity and has been associated with trastuzumab resistance while retaining sensitivity to lapatinib (25, 26, 60–63).

**Figure 2.**
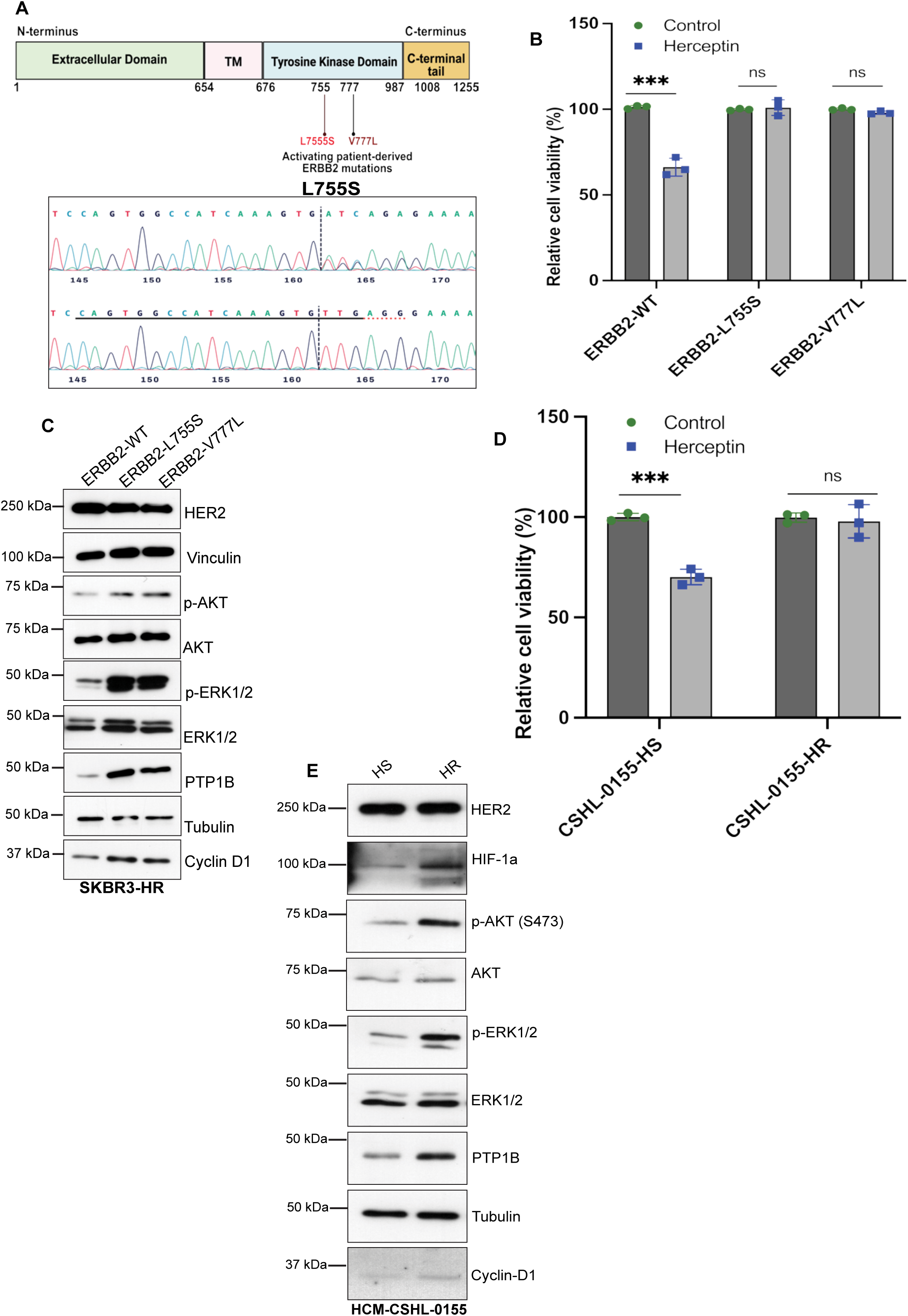
PTP1B was elevated in ERBB2-mutant and trastuzumab-resistant HER2-positive tumor organoid models. **(A)** Top: Schematic of the HER2 (ERBB2) receptor showing the extracellular, transmembrane (TM), intracellular tyrosine kinase, and C-terminal domains. The clinically relevant activating mutations L755S and V777L are located within the kinase domain, as shown. Created in BioRender. M, V. (2026) https://BioRender.com/sw1qf8n. **Bottom:** Sanger sequencing chromatograms confirming successful CRISPR–Cas9-mediated introduction of the endogenous ERBB2 L755S and V777L (shown in Figure S2A) mutations into SKBR3 cells. **(B)** Cell viability measured by the CellTiter-Glo (CTG) assay in SKBR3 cells expressing wild-type (WT) ERBB2 or the endogenous L755S and V777L mutant variants following trastuzumab treatment (72 h), showing no sensitivity of the mutant cells to trastuzumab compared with WT cells. **(C)** Immunoblot analysis of PTP1B and indicated signaling proteins in SKBR3 cells expressing WT or mutant ERBB2 (L755S and V777L), showing increased PTP1B expression, elevated HIF-1α levels, sustained p-AKT and p-ERK signaling, and increased Cyclin D1 expression in mutant cells. **(D)** CellTiter-Glo (CTG) assay showing the response of HCM-CSHL-0155-HS and HCM-CSHL-0155-HR tumor organoids to trastuzumab treatment (72 h). HR organoids exhibited reduced sensitivity to trastuzumab. **(E)** Immunoblot analysis of PTP1B, HIF-1α, and indicated signaling proteins in HS and HR tumor organoids, demonstrating increased PTP1B expression, elevated HIF-1α levels, sustained p-AKT and p-ERK signaling, and increased Cyclin D1 expression in resistance organoids. Data are presented as mean ± SD from three independent biological experiments unless otherwise indicated. Data were normalized to the respective control for visualization. Statistical significance in B and E was determined using two-way ANOVA followed by Bonferroni’s multiple comparisons test. ns, not significant; *P < 0.05; **P < 0.01; ***P < 0.001; ****P < 0.0001.

To determine whether these clinically relevant *ERBB2* alterations recapitulate the effects on signaling observed in other resistance models, we introduced L755S and V777L mutations into the endogenous *ERBB2* locus of SKBR3 cells using CRISPR–Cas9-mediated editing (**Figure 2A; Supplementary Figure S2A**). Both mutant models were resistant to trastuzumab; L755S cells were also resistant to lapatinib, whereas V777L cells retained sensitivity to lapatinib (**Figure 2B; Supplementary Figure S2B**). Both mutant cell lines exhibited increased PTP1B expression, AKT and ERK phosphorylation, and elevated Cyclin D1 and HIF-1α expression relative to wild-type cells (**Figure 2C**). We also generated a trastuzumab-resistant derivative of the HER2-positive patient-derived organoid line HCM-CSHL-0155-HR through prolonged trastuzumab exposure (**Figure 2D, Supplementary Figure S2C**). Resistant organoids displayed increased PTP1B expression, increased AKT and ERK phosphorylation, and elevated Cyclin D1 expression (**Figure 2E**). Moreover, the *ERBB2* L755S and V777L-mutant cells, trastuzumab-resistant organoids, and SKBR3-HR orthotopic xenografts exhibited increased growth relative to their respective trastuzumab-sensitive controls (**Supplementary Figures S2D–S2H**).

Collectively, these findings demonstrate that increased PTP1B expression accompanied trastuzumab resistance in several distinct models. These findings associate PTP1B upregulation with proliferative and survival-associated signaling across multiple trastuzumab-resistance models and provide the rationale to determine whether PTP1B is functionally required for maintenance of the resistance phenotype and whether its inhibition can restore trastuzumab sensitivity.

### Pharmacological inhibition of PTP1B suppressed growth of trastuzumab-resistant models

Trastuzumab treatment inhibited the growth of sensitive SKBR3 and BT474 cells (**Figures 3A, 3D** and **Supplementary Figure S3A-B**) but had little effect in the resistance models (**Figures 3B, C, E-G and Supplementary Figures S3A, S3C, S3D–S3F**). The small-molecule PTP1B inhibitor DPM-1003 (64) suppressed cell growth in intrinsic resistance cell models, including JIMT1 and HCC1569, acquired resistance cell lines SKBR3-HR and BT474-HR, and cells harboring the endogenous ERBB2 kinase domain mutations L755S and V777L (**Figures 3A-3G and Supplementary Figures S3A-S3F**). These results identify a pharmacological vulnerability across the resistance models tested.

**Figure 3.**
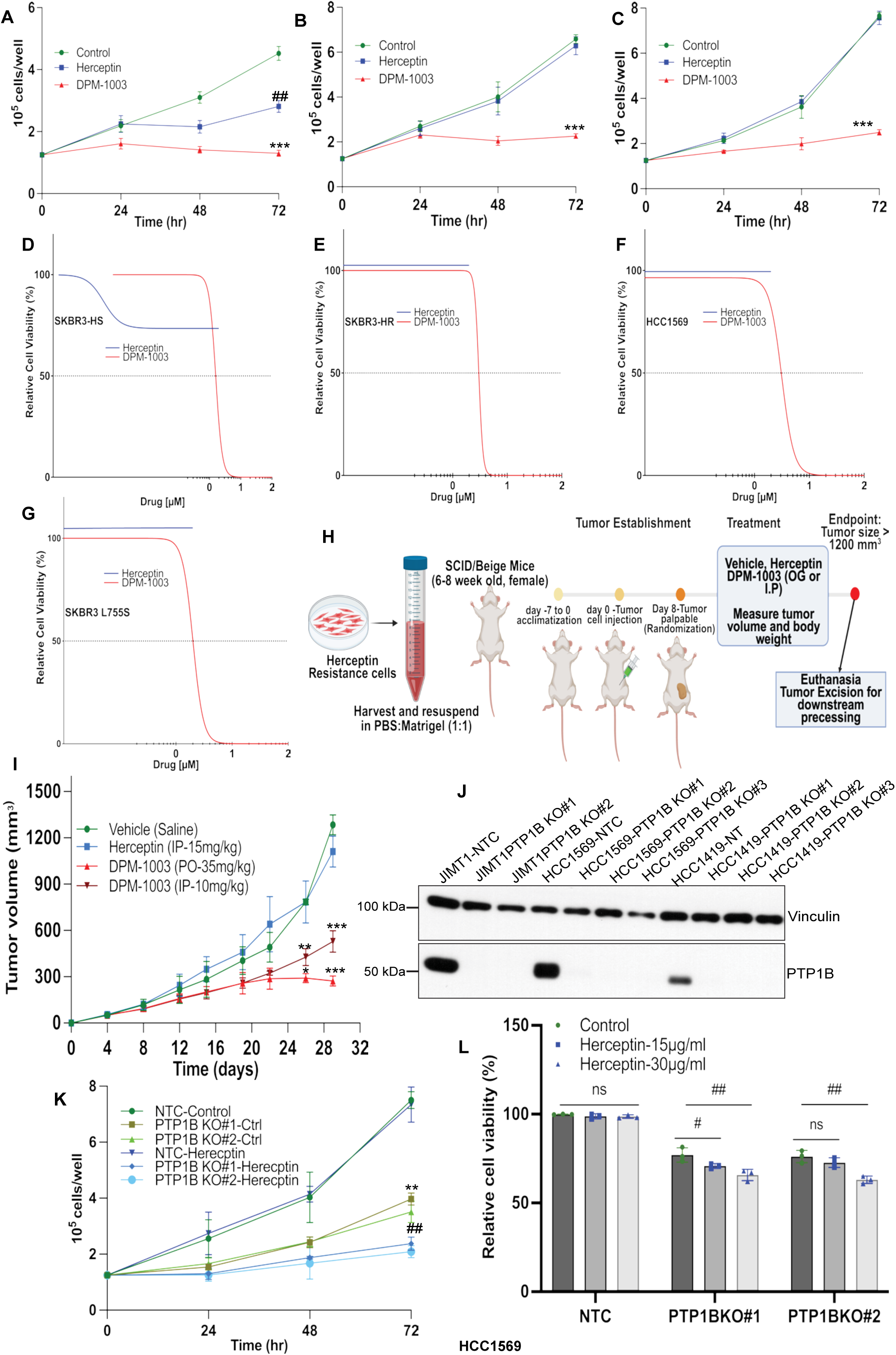
Pharmacological and genetic inhibition of PTP1B overcame trastuzumab resistance. **(A–C)** Cell proliferation measured by hemocytometer counting in SKBR3-HS **(A)**, SKBR3-HR **(B)**, and HCC1569 **(C)** cells following treatment with vehicle, trastuzumab (32 μg/mL), or the PTP1B inhibitor DPM-1003 (2 μM) for 72 h. DPM-1003 suppressed proliferation in trastuzumab-resistant cells, whereas trastuzumab had little effect. **(D–G)** Cell viability was measured by the CellTiter-Glo (CTG) assay following 72 h of dose-dependent treatment with trastuzumab or the PTP1B inhibitor DPM-1003 in SKBR3-HS **(D)**, SKBR3-HR **(E)**, HCC1569 **(F)**, and SKBR3 cells expressing the endogenous ERBB2 L755S mutation **(G),** demonstrating that PTP1B inhibition markedly reduced growth in all cell models, whereas trastuzumab showed inhibition only in sensitive cells. **(H)** Schematic illustration of the orthotopic mammary fat pad xenograft experimental design in SCID/Beige mice implanted with JIMT1 cells and treated with vehicle, trastuzumab, or DPM-1003. Created in BioRender. M, V. (2026) https://BioRender.com/sw1qf8n **(I)** Tumor growth curves of JIMT1 xenografts treated with vehicle, trastuzumab (15 mg/kg), or DPM-1003 (35 mg/kg, PO; 10 mg/kg, intraperitoneal), administered twice weekly. **(J)** Immunoblot analysis confirming CRISPR–Cas9-mediated knockout of PTP1B in the intrinsic trastuzumab-resistant cell lines JIMT1, HCC1419, and HCC1569. **(K–L)** Cell proliferation **(K)** and Cell Titer-Glo (CTG) cell viability **(L)** assays of HCC1569 cells expressing a non-targeting control (NTC) or *PTPN1* knockout following trastuzumab treatment (72 h), demonstrating restoration of trastuzumab sensitivity following *PTPN1* loss. Data are presented as mean ± SD from three independent biological experiments unless otherwise indicated. Data were normalized to the respective control for visualization. Statistical significance was determined by two-way ANOVA (L) followed by Bonferroni’s multiple-comparisons test or by mixed-effects analysis (A–C, I, and K) followed by Tukey’s multiple-comparisons test. Asterisks (*, **, ***) indicate comparisons between control and DPM-1003-treated groups or between untreated control and PTP1B-KO groups; number signs (#, ##, ###) indicate comparisons between control and Herceptin-treated groups or between untreated PTP1B-KO and PTP1B-KO + Herceptin-treated groups. Tumor growth (I) was evaluated in three independent cohorts, with four mice per treatment group in each cohort (total n = 12 mice per group). */#P < 0.05; **/##P < 0.01; ***/###P < 0.001.

We established orthotopic mammary fat pad xenografts using the intrinsically resistant JIMT1 cell line; consistent with the *in vitro* findings, trastuzumab did not affect tumor growth, whereas DPM-1003 suppressed tumor progression (**Figures 3H-3I**). Similarly, DPM-1003 inhibited the growth of trastuzumab-resistant HER2-positive tumor organoids, whereas trastuzumab exhibited no activity (**Supplementary Figures S3G-S3H**). Collectively, these experiments establish that pharmacological inhibition of PTP1B is active against trastuzumab-resistant tumors and organoids.

### Genetic ablation of PTP1B restored trastuzumab responsiveness in selected resistance models

Having established that pharmacological inhibition of PTP1B suppressed the growth of trastuzumab-resistant models, we tested whether genetic ablation of *PTPN1* similarly restores trastuzumab sensitivity. We deleted *PTPN1* using CRISPR–Cas9 in multiple resistant HER2-positive cell models (**Figures 3J, 4A & 4D**). Deletion reduced growth and restored a response to trastuzumab in both intrinsic-resistance HCC1569 and HCC1419 cells and in acquired-resistance SKBR3-HR and BT474-HR cells (**Figures 3K-L, 4B-C, 4E–4F, Supplementary Figures S4A**). In JIMT1 cells, deletion reduced growth without restoring trastuzumab responsiveness (**Supplementary Figures S4B-S4C**). Therefore, PTP1B supports trastuzumab-resistant cell growth, but its contribution to responsiveness varies between models.

Trastuzumab-resistant cells expressing elevated PTP1B exhibited lower phosphotyrosine levels than sensitive controls, whereas CRISPR-mediated deletion of PTP1B resulted in the expected increase in phosphotyrosine abundance in both intrinsic and acquired resistance models (**Fig. 4G–4I and Supplementary Figures S4D–S4E**). This response prompted quantitative profiling of PTP1B-dependent phosphorylation events and signaling pathways. SKBR3-HR cells were selected for phosphoproteomic analysis. HCC1569 and additional resistance models were used to assess the reproducibility of key biochemical and functional findings.

**Figure 4.**
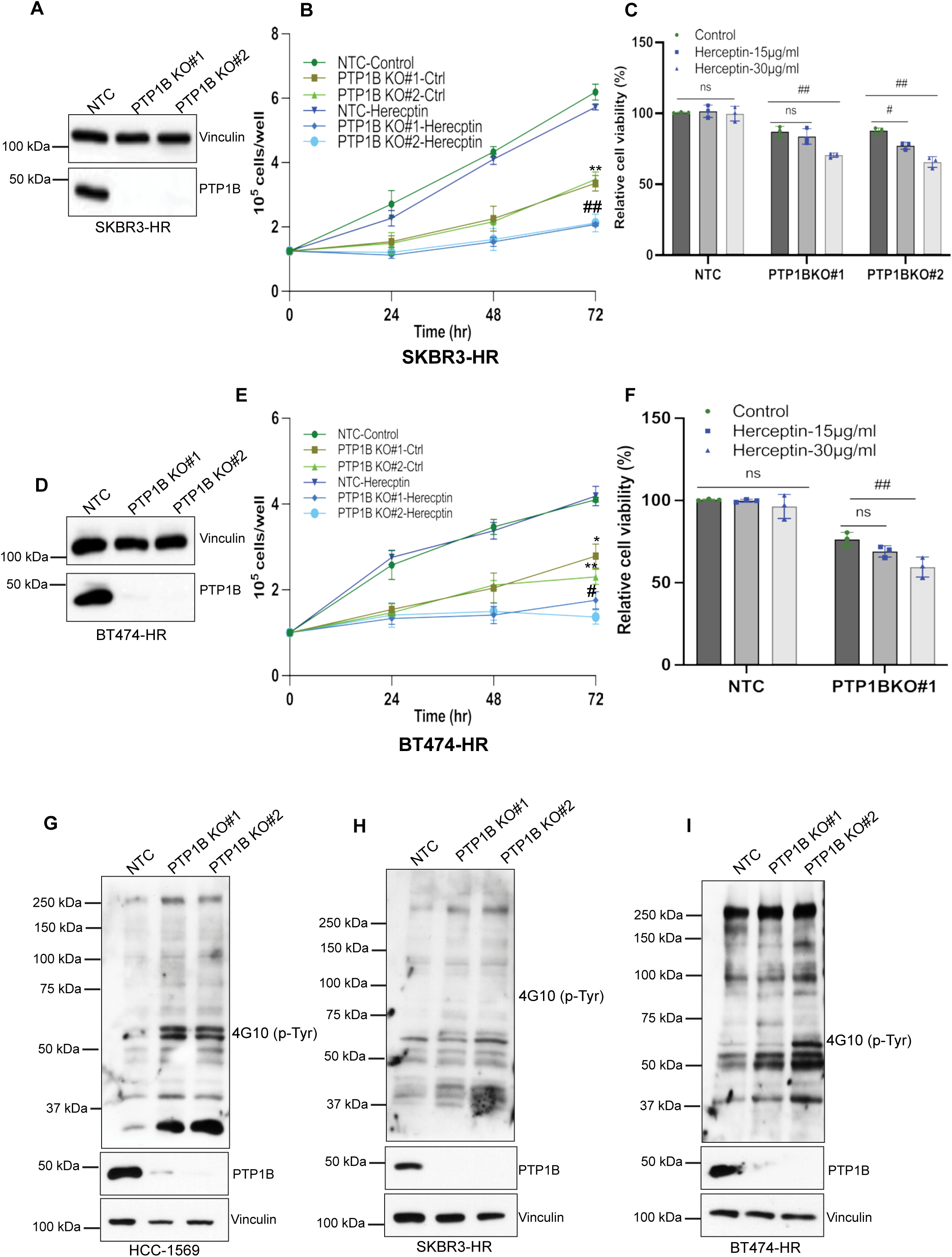
Genetic ablation of PTP1B restored trastuzumab sensitivity in selected Herceptin-resistant cell models and increased phosphotyrosine signaling. **(A)** Immunoblot analysis confirming CRISPR–Cas9-mediated knockout of *PTPN1* in acquired trastuzumab-resistant SKBR3-HR cells using independent sgRNAs. **(B–C)** Cell proliferation (B; hemocytometer counting) and cell viability (C; CTG assay) of SKBR3-HR cells expressing a non-targeting control (NTC) or *PTP1N1* knockout following trastuzumab treatment (72 h), demonstrating restoration of trastuzumab sensitivity upon *PTPN1* loss. **(D)** Immunoblot analysis confirming CRISPR–Cas9-mediated knockout of *PTPN1* in acquired trastuzumab-resistant BT474-HR cells using independent sgRNAs. **(E–F)** Cell proliferation (E; hemocytometer counting) and cell viability (F; CTG assay) of BT474-HR cells expressing NTC or *PTPN1* knockout following trastuzumab treatment (72 h), demonstrating restoration of trastuzumab sensitivity upon PTP1B loss. **(G–I)** Immunoblot analysis of global phosphotyrosine (p-Tyr) levels in control and *PTPN1* knockout cells in the intrinsic trastuzumab-resistantHCC1569 cells **(G)** and acquired trastuzumab-resistant SKBR3-HR cells **(H)** and BT474-HR **(I)** cells, demonstrating increased phosphotyrosine signaling following PTP1B loss. Data are presented as mean ± SD from three independent biological experiments unless otherwise indicated. Data were normalized to the respective control for visualization. Statistical significance was determined by two-way ANOVA (C and F), followed by Bonferroni’s multiple-comparisons test, or by mixed-effects analysis (B and F), followed by Tukey’s multiple-comparisons test. Asterisks (*, **, ***) indicate comparisons between control and DPM-1003-treated groups or between untreated control and PTP1B-KO groups; number signs (#, ##, ###) indicate comparisons between control and Herceptin-treated groups or between untreated PTP1B-KO and PTP1B-KO + Herceptin-treated groups. */#P < 0.05; **/##P < 0.01; ***/###P < 0.001.

### Phosphotyrosine profiling and substrate trapping identified CFL1 and HSP90α as PTP1B substrates

Phosphotyrosine enrichment and immunoblotting confirmed increased phosphorylation in SKBR3-HR cells following PTP1B deletion (**Figure 5A**). We then performed TMT-based quantitative phosphoproteomics using control cells and cells generated with two independent *PTPN1*-targeting sgRNAs (**Supplementary Figure S5A**).

**Figure 5.**
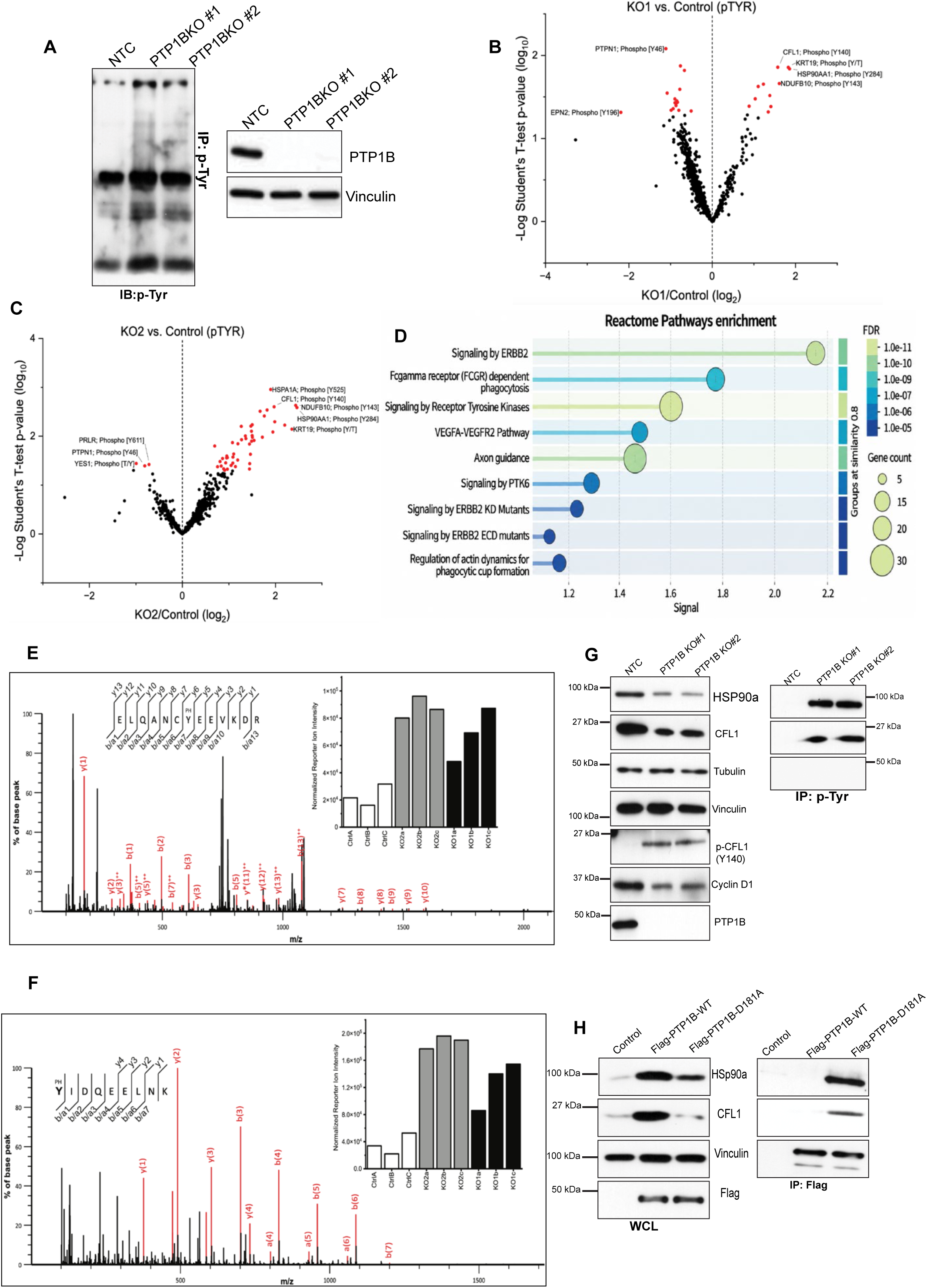
Identification of CFL1 and HSP90α as direct substrates of PTP1B in trastuzumab-resistant cells. **(A)** Enrichment of phosphotyrosine (p-Tyr) proteins from SKBR3-HR cells expressing a non-targeting control (NTC) or *PTPN1* knockout (two independent sgRNAs) using p-Tyr-1000 magnetic immunoaffinity beads, followed by immunoblotting with an anti-phosphotyrosine antibody, showing increased phosphotyrosine levels following *PTPN1* loss. **(B–C)** Volcano plots of differentially phosphorylated phosphotyrosine peptides comparing NTC and *PTPN1* knockout cells generated with two independent sgRNAs (C, KO#1; D, KO#2). Significantly regulated phosphotyrosine sites (*P* < 0.05) are highlighted in red. **(D)** Reactome pathway enrichment analysis of differentially regulated phosphotyrosine proteins showing enrichment of HER2/ERBB2 signaling, receptor tyrosine kinase signaling, constitutive PI3K signaling, HIF signaling, and tyrosine kinase inhibitor resistance pathways (Sup. Fig. S6C-D). **(E–F)** Representative MS/MS spectra of phosphopeptides from CFL1 (E) and HSP90a (F), with annotated fragment ions supporting identification. Insets show the intensity of the TMT reporter ions used for relative quantification. **(G)** Immunoprecipitation of phosphotyrosine-containing proteins followed by immunoblotting for CFL1 and HSP90α1, confirming increased tyrosine phosphorylation of both proteins in *PTPN1* knockout cells (right). Corresponding whole-cell lysates are shown (left). **(H)** Substrate-trapping assay using FLAG-tagged PTP1B wild-type (WT) or the catalytically inactive D181A mutant, demonstrating preferential interaction of CFL1 and HSP90α1 with the trapping mutant, consistent with direct substrate binding (right). Corresponding whole-cell lysates are shown (left). Quantification from three independent biological experiments is presented in Sup Figure S6E-F. Data are presented as mean ± SD from three independent biological experiments unless otherwise indicated. Volcano plots are shown for the indicated experimental conditions. Data points represent p-Tyr sites that met the filtering criteria in individual biological replicates. Red data points indicate p-Tyr sites that met the filtering criteria in ≥3 biological replicates, with a fold change ≥1 and P ≤ 0.05.

Across enriched fractions, 26,045 peptides were identified (False Discovery Rate (FDR) <1%), of which 20,867 were quantified, including 6,084 phosphorylated peptides. The analysis included 786 peptides with localized tyrosine phosphorylation or ambiguously localized phosphorylation in a tyrosine-containing sequence (**Supplementary Tables S1–S3**). Differential analysis identified changes in phosphorylation following PTP1B deletion (**Figures 5B-5C; Supplementary Figure S5B**). Functional enrichment analysis implicated receptor tyrosine kinase signaling, HER2, PI3K signaling, and HIF-associated pathways (**Figure D; Supplementary Figures S5C-S5D**). These findings are consistent with our earlier biochemical observations that trastuzumab-resistant models exhibit elevated PTP1B expression, sustained p–AKT and p-ERK signaling, and increased HIF-1α levels (**Figures 1** and **2**), providing independent phosphoproteomic support for PTP1B-dependent signaling associated with trastuzumab resistance.

Cofilin-1 (CFL1) and HSP90α were selected for validation because their phosphopeptide signals increased after PTP1B deletion (**Figures 5E-5F**). Immunoblotting using phospho-specific antibodies detected increased phosphorylation of CFL1 (Y140) in whole-cell lysates following PTP1B deletion (**Figure 5G**). Due to the lack of a phospho-specific antibody for HSP90α (Y284), phosphotyrosine immunoprecipitation followed by immunoblotting confirmed increased endogenous tyrosine phosphorylation of both CFL1 and HSP90α in PTP1B-deficient cells (**Figure 5G**). To determine whether these proteins are direct PTP1B substrates, we performed substrate-trapping assays (45, 49) using the catalytically attenuated PTP1B D181A mutant. The substrate-trapping mutant, but not wild-type PTP1B, formed stable complexes with both CFL1 and HSP90α (**Figure 5H; Supplementary Fig S5E-F**). Collectively, these findings identify CFL1 and HSP90α as direct PTP1B substrates.

### PTP1B regulated CFL1 and HSP90α abundance and proteasome-dependent turnover

Immunoblot analysis revealed that the abundance of both CFL1 and HSP90α was increased in intrinsic and acquired trastuzumab-resistant models relative to trastuzumab-sensitive controls, in parallel with increased PTP1B expression (**Figures 6A and 6B**). Similar increases were observed in trastuzumab-resistant tumor organoids and orthotopic xenografts (**Figures 6C** and **6D).** Previous studies have shown that HSP90α stabilizes HIF-1α under cellular stress and HIF-1α acts downstream of HER2 signaling (15, 16, 65–69), whereas CFL1 has been implicated in tumor cell survival, cytoskeletal regulation, and apoptosis-associated processes (70–72). Consistent with these reports and our earlier observations (Figures 1 and 2), resistance models with elevated HSP90α also exhibited increased HIF-1α expression, whereas increased CFL1 expression was accompanied by elevated Bcl2 levels (**Figures 6A-D)**. Together, these observations are consistent with a survival-associated signaling program in trastuzumab-resistant cells. These associations prompted us to test whether PTP1B regulated substrate abundance.

**Figure 6.**
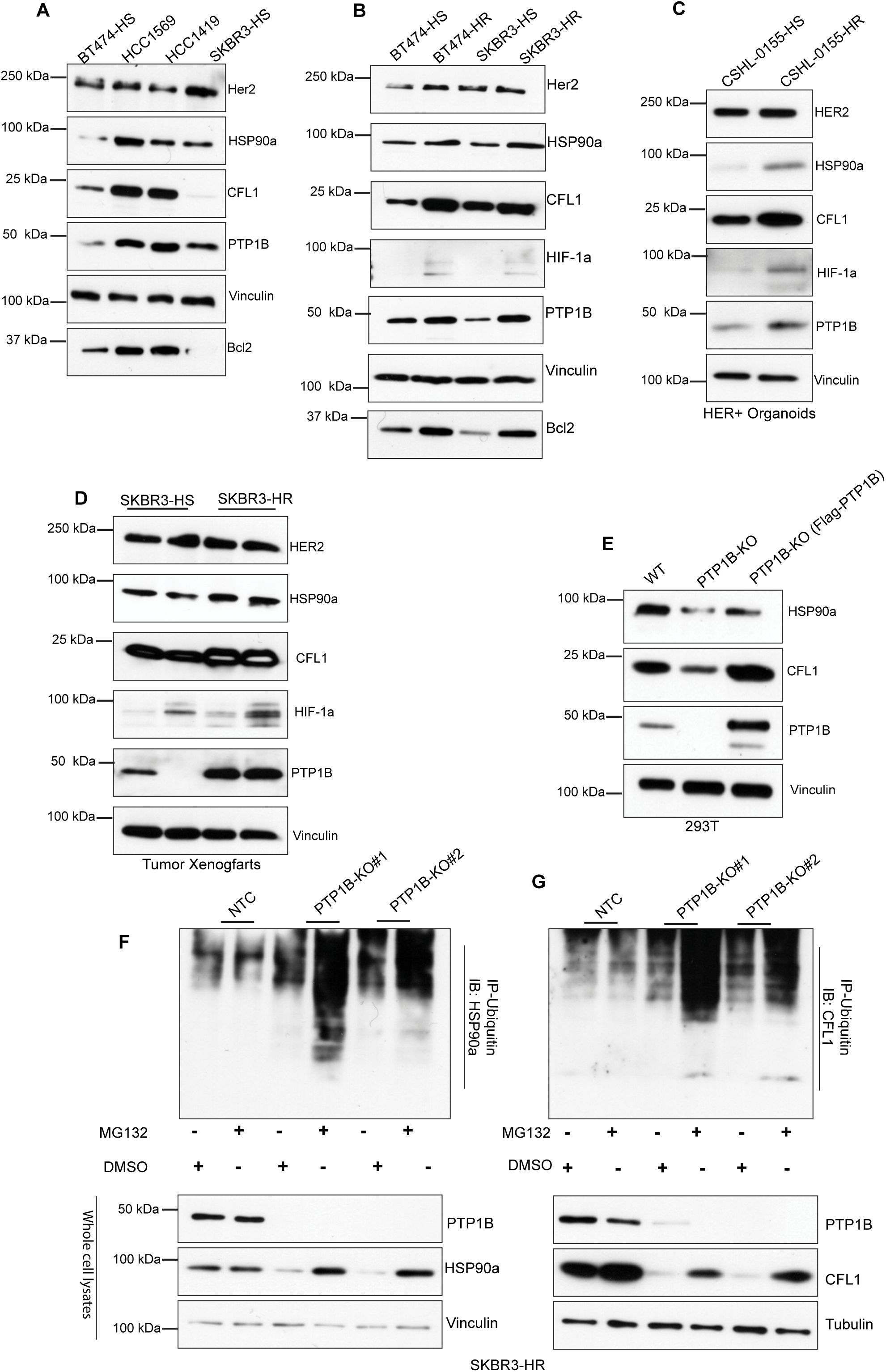
Levels of CFL1 and HSP90α were elevated in trastuzumab-resistance models and were regulated by PTP1B-dependent proteasomal degradation. **(A–B)** Immunoblot analysis of CFL1, HSP90α1, PTP1B, HIF-1α, and indicated signaling proteins in trastuzumab-sensitive (HS; BT474 and SKBR3) and intrinsic trastuzumab-resistance (**A**; HCC1569 and HCC1419) or acquired trastuzumab-resistance (**B**; SKBR3-HR and BT474-HR) HER2-positive breast cancer cell lines, showing upregulation of PTP1B, CFL1, HSP90α, HIF-1α, and downstream p-AKT/p-ERK levels in resistance models. **(C–D)** Immunoblot analysis of CFL1, HSP90α, PTP1B, HIF-1α, and the indicated signaling proteins in HER2-positive tumor organoids (HCM-CSHL-0155; **C**) and SKBR3 xenograft tumors (**D**), showing elevated expression in trastuzumab-resistant models compared with sensitive controls. **(E)** Immunoblot analysis of PTP1B-null HEK293T cells following re-expression of PTP1B, demonstrating restoration of CFL1 and HSP90α protein levels. **(F–G)** Analysis of HSP90α1 (**F**) and CFL1 (**G**) ubiquitination and proteasomal degradation. HA-ubiquitin pull-down assays in SKBR3-HR control and ***PTPN1*** knockout cells treated with MG132 (5 μM) demonstrate increased ubiquitination of HSP90α1 and CFL1 following PTP1B loss. Immunoblot analysis of corresponding whole-cell lysates shows accumulation of HSP90α1 and CFL1 following MG132 treatment, consistent with proteasome-dependent degradation (bottom). Data are presented from three independent biological experiments unless otherwise indicated.

Loss of PTP1B reduced the abundance of both CFL1 and HSP90α, whereas re-expression of PTP1B restored their protein levels in HEK293T cells (**Figure 6E**). Because both proteins exhibited increased tyrosine phosphorylation following PTP1B deletion (**Figure 5**), and tyrosine phosphorylation has been implicated in regulating CFL1 turnover and HSP90α function (73–77), we investigated whether PTP1B influences their stability through the ubiquitin–proteasome pathway. In PTP1B-deficient SKBR3-HR cells, CFL1 and HSP90α protein levels were restored by treatment with the proteasome inhibitor MG132 (**Figures 6F-6G**), indicating that their degradation is proteasome-dependent. In addition, ubiquitination pull-down assays demonstrated increased ubiquitination and degradation of both proteins following PTP1B deletion (**Figures 6F-6G**). Together, these findings support increased ubiquitin-associated, proteasome-dependent degradation of CFL1 and HSP90α after PTP1B loss. Combined with the phosphoproteomic and substrate-trapping analyses, these findings demonstrate that PTP1B-dependent dephosphorylation stabilizes CFL1 and HSP90α by limiting their ubiquitination and proteasomal degradation.

### PTP1B targeting suppressed survival signaling and induced apoptotic cell death

CRISPR–Cas9-mediated deletion of *PTPN1* reduced CFL1 and HSP90α abundance and attenuated AKT and ERK phosphorylation in acquired-resistance SKBR3-HR, intrinsically resistant HCC1569, and ERBB2-L755S mutant cells (**Figures 7A–7C**). Given the established roles of HSP90α in regulating HIF-1α stability and HER2-associated signaling (15, 16, 65–67, 77–79), we also examined HIF-1α and downstream survival– and proliferation-associated proteins. PTP1B deletion reduced HIF-1α, Bcl2, and Cyclin D1 expression, accompanied by increased cleavage of PARP and caspase-3, consistent with induction of apoptosis (**Figures 7A–7C**). These findings indicate that PTP1B loss destabilizes CFL1 and HSP90α, suppresses oncogenic and stress-adaptive signaling, and activates apoptotic pathways across distinct trastuzumab-resistance models.

**Figure 7.**
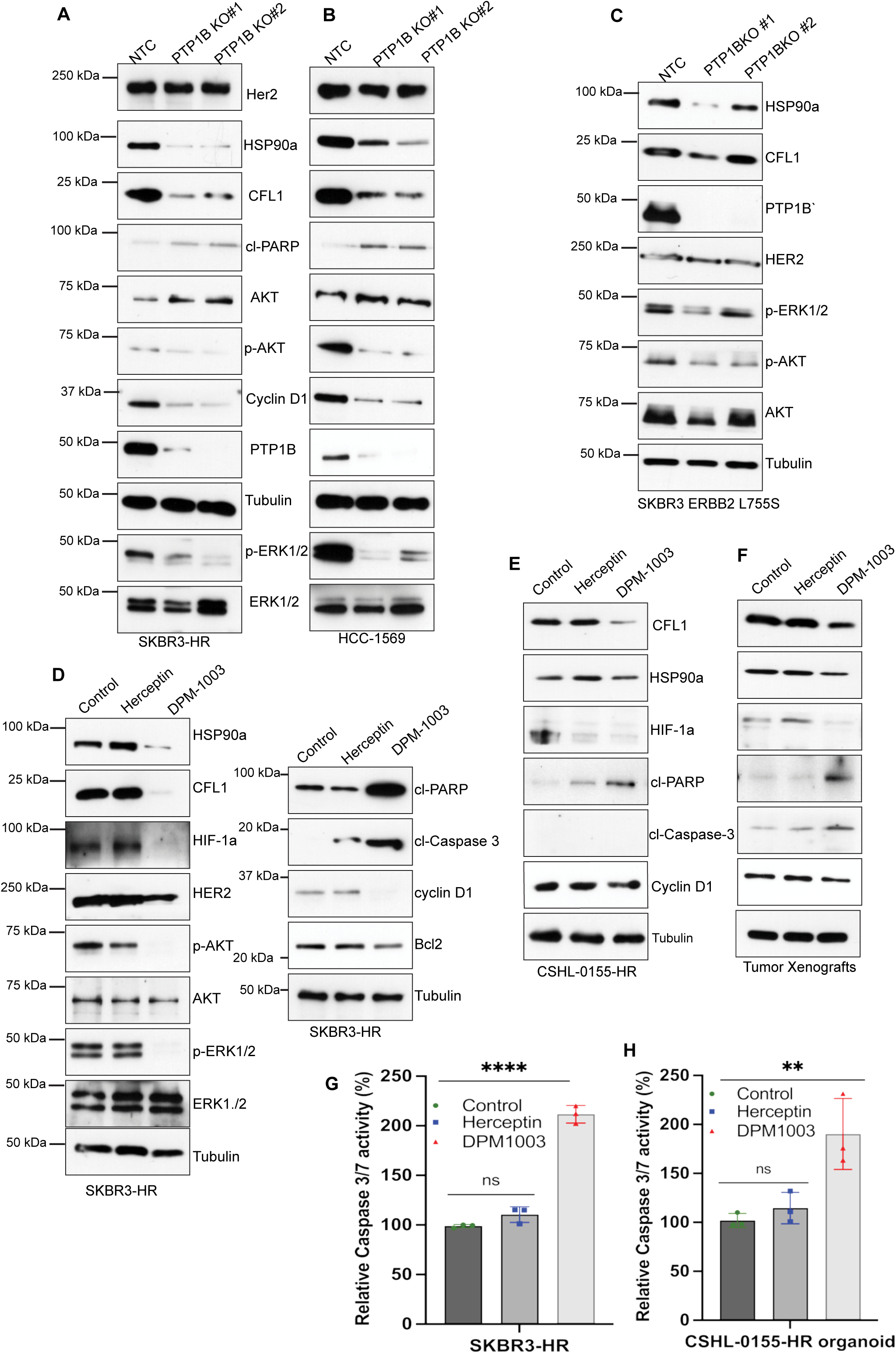
PTP1B inhibition disrupted oncogenic and stress-adaptive signaling and induced apoptosis in trastuzumab-resistant cells. **(A–C)** Immunoblot analysis following CRISPR–Cas9-mediated knockout of PTP1B in acquired trastuzumab-resistant SKBR3-HR **(A)**, intrinsic trastuzumab-resistant HCC1569 **(B)**, and SKBR3 cells expressing the endogenous ERBB2 L755S mutation **(C)**, showing reduced CFL1 and HSP90α expression together with decreased p-AKT and p-ERK levels. **(D)** Immunoblot analysis of SKBR3-HR cells treated with vehicle, trastuzumab (32 μg/mL), or the PTP1B inhibitor DPM-1003 (2 μM) for 48 h. DPM-1003, but not trastuzumab, reduced CFL1, HSP90α, HIF-1α, phospho-AKT, phospho-ERK1/2, Bcl2, and Cyclin D1 levels while increasing cleaved PARP and cleaved caspase-3, consistent with induction of apoptosis. **(E–F)** Immunoblot analysis of HER2-positive trastuzumab-resistant tumor organoids, HCM-CSHL-0155-HR **(E),** and orthotopic mammary fat pad xenografts **(F)** treated with vehicle, trastuzumab, or DPM-1003, showing reduced CFL1, HSP90α, HIF-1α, Bcl2, and Cyclin D1 expression together with increased apoptotic markers following PTP1B inhibition. **(G–H)** Caspase-3/7 activity assays in SKBR3-HR cells **(G)** and HCM-CSHL-0155 trastuzumab-resistant tumor organoids **(H)** treated with vehicle, trastuzumab, or DPM-1003, demonstrating increased apoptotic activity following PTP1B inhibition. Data are presented as mean ± SD from three independent biological experiments unless otherwise indicated. Data were normalized to the respective control for visualization. Statistical significance in G and H was determined by one-way ANOVA followed by Bonferroni’s multiple-comparisons test. ns, not significant; *P < 0.05; **P < 0.01; ***P < 0.001, ****P < 0.0001.

Similar to genetic deletion, DPM-1003 treatment reduced CFL1 and HSP90α abundance, attenuated AKT and ERK phosphorylation, and decreased HIF-1α, Bcl2, and Cyclin D1 expression, accompanied by increased PARP and caspase-3 cleavage in different trastuzumab-resistant cell models (**Figure 7D**; **Supplementary Figures S7A-7C**). Moreover, trastuzumab-resistant tumor organoids and orthotopic xenografts also showed decreased CFL1, HSP90α, and HIF-1α abundance and increased apoptotic markers (**Figures 7E and 7F**). Furthermore, functional assays confirmed increased caspase-3/7 activity following PTP1B inhibition in resistant cells and tumor organoids (**Figures 7G and 7H, Supplementary Figures S7D and S7E**). Together, these results demonstrate that genetic and pharmacological inhibition of PTP1B produced concordant molecular and apoptotic responses, supporting a role for PTP1B in maintaining proliferative and survival-associated signaling during trastuzumab resistance.

### HSP90 inhibition also reduced viability of trastuzumab-resistant cells

Treatment with the PTP1B inhibitor DPM-1003 or the HSP90 inhibitor SNX-5422 (80) decreased ATP-based viability signals in SKBR3-HR, ERBB2-L755S mutant, and HCC1569 cells, whereas trastuzumab alone had little effect (**Figures 8A–8C**). These findings support an HSP90-associated vulnerability in trastuzumab-resistant cells.

**Figure 8.**
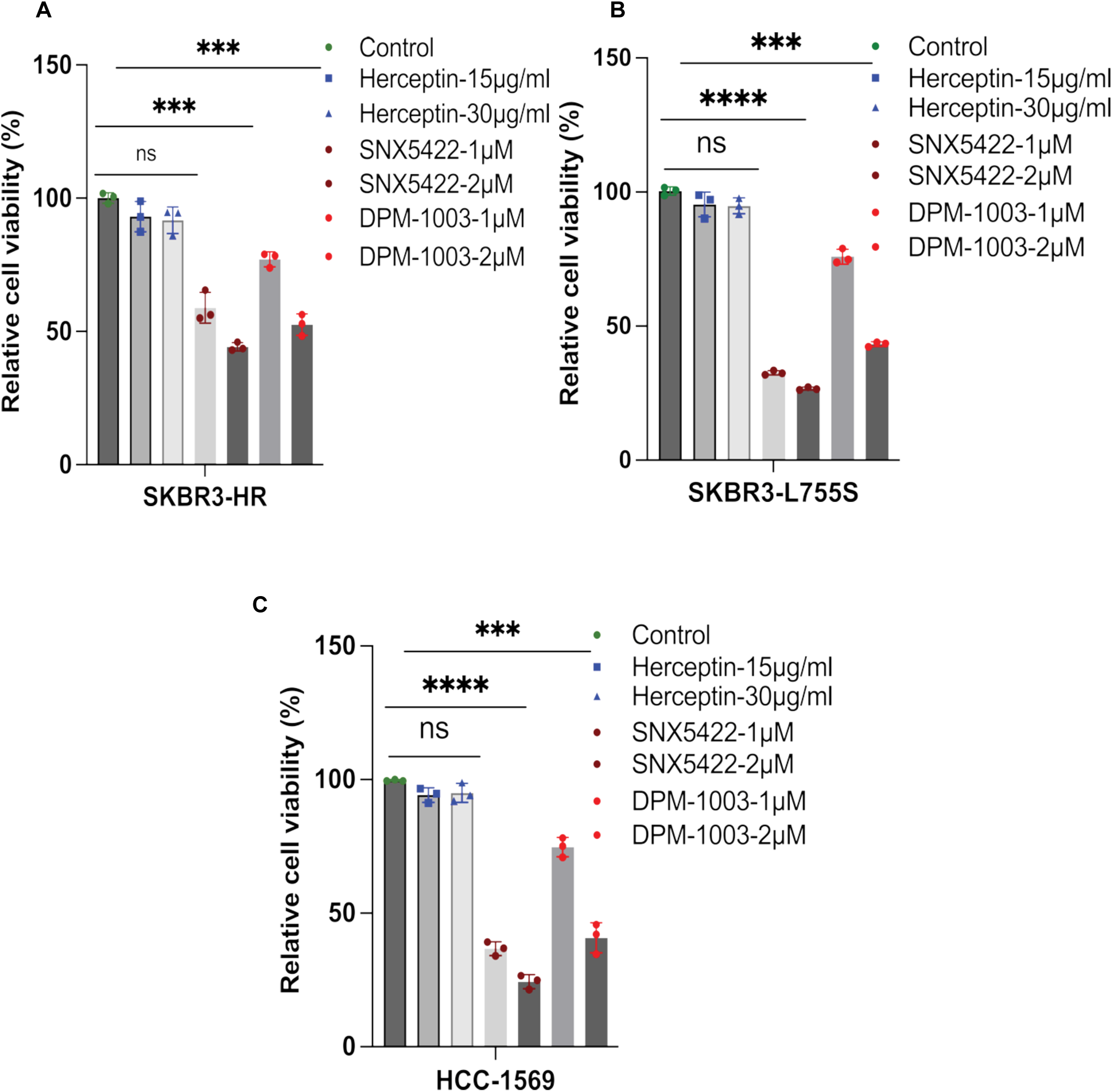
Targeting PTP1B or HSP90α overcame trastuzumab resistance in Herceptin resistance cell models. **(A)** Cell viability measured by the CellTiter-Glo (CTG) assay in SKBR3-HR cells treated with vehicle, trastuzumab, the PTP1B inhibitor DPM-1003 (PTP1Bi), or the HSP90α inhibitor SNX-5422 (HSP90αi) for 72 h. Pharmacological inhibition of PTP1B or HSP90α significantly reduced cell viability, whereas trastuzumab showed no inhibition. **(B–C)** Cell viability was measured by the CellTiter-Glo (CTG) assay in SKBR3 cells expressing the endogenous ERBB2 L755S mutation **(B)** and intrinsic trastuzumab-resistant HCC1569 cells **(C)** following treatment with vehicle, trastuzumab, DPM-1003, or SNX-5422 for 72 h. Inhibition of PTP1B or HSP90α significantly reduced cell viability, whereas trastuzumab showed limited activity. Data are presented as mean ± SD from three independent biological experiments unless otherwise indicated. Data were normalized to the respective control for visualization. Statistical significance in A, B, and C was determined using one-way ANOVA followed by Bonferroni’s multiple comparisons test. ns, not significant; *P < 0.05; **P < 0.01; ***P < 0.001, ****P < 0.0001.

Together, our findings identify PTP1B as a therapeutic vulnerability in trastuzumab-resistant HER2-positive breast cancer models. PTP1B targeting reduced CFL1 and HSP90 abundance, suppressed survival-associated signaling, and induced apoptosis. These results support a model linking PTP1B-dependent regulation of substrate stability to trastuzumab-resistant cell survival, although the causal contributions of CFL1 and HSP90 remain to be established.

## Discussion

Although trastuzumab has substantially improved patient outcomes in HER2-positive breast cancer, intrinsic and acquired resistance limit its long-term clinical benefit (13, 17). In this study, we identify PTP1B as a critical regulator of trastuzumab resistance across multiple HER2-positive breast cancer models, including intrinsic– and acquired-resistance cells, *ERBB2*-mutant models, tumor organoids, and orthotopic xenografts. PTP1B expression was consistently elevated in resistance models and was associated with sustained PI3K–AKT/ERK signaling, increased Cyclin D1 expression, and elevated HIF-1α levels (**Figures 1 and 2**). In contrast to the predominantly cytostatic effects of trastuzumab in sensitive cells (56–59), genetic ablation or pharmacological inhibition of PTP1B with DPM-1003 overcame trastuzumab resistance in multiple independent models, inducing apoptosis and suppressing tumor growth (**Figures 3 and 7**). In addition, PTP1B deletion reduced growth and restored trastuzumab responsiveness in selected models. These findings extend previous studies establishing a tumor-promoting role for PTP1B in HER2-driven breast cancer (39–41) and demonstrate that PTP1B is functionally required for maintenance of the resistance phenotype in a subset of HER2-positive breast cancer models. Nevertheless, growth dependency and restoration of trastuzumab responsiveness were distinct outcomes. JIMT1 cells remained resistant to trastuzumab after PTP1B deletion despite reduced growth, indicating that PTP1B-independent mechanisms can maintain resistance. Moreover, DPM-1003 monotherapy activity establishes susceptibility to PTP1B targeting, not pharmacological resensitization to trastuzumab. Combination experiments will be needed to determine whether acute PTP1B inhibition restores trastuzumab responses and whether the combination provides benefit beyond either intervention alone.

Our phosphoproteomic analyses implicated signaling networks relevant to HER2 biology and drug resistance, consistent with previous studies of resistance-associated changes in protein phosphorylation (14–16,18, 50,61,81–83). This provides independent support that PTP1B regulates specific phosphotyrosine-dependent signaling events associated with trastuzumab resistance, rather than supporting widespread or indiscriminate phosphotyrosine regulation. A mechanistic advance of our study is the identification of CFL1 and HSP90α as direct substrates of PTP1B, the abundance of which were decreased after PTP1B loss. Increased phosphorylation, preferential association with the substrate-trapping mutant over wild-type phosphatase, restoration of abundance by PTP1B re-expression, and MG132-sensitive turnover support a connection between PTP1B activity and substrate stability. This expands the mechanisms through which PTP1B can support oncogenic signaling, beyond previously characterized substrates such as SRC, p62Dok, CDK3, BCAR3, BCAR1, CDCP1, and RNF213 (41–55), highlighting the diverse mechanisms by which PTP1B-dependent tyrosine dephosphorylation regulates oncogenic and stress-adaptive signaling. Our findings are consistent with previous evidence that tyrosine phosphorylation of CFL1 can promote its ubiquitination and proteasomal degradation (74,76) and that phosphorylation of HSP90α regulates its chaperone function and can influence its stability and degradation (75, 77). Nevertheless, phosphosite substitution, protein half-life determination, and rescue with wild-type versus catalytically inactive PTP1B will help define the causal relationship between phosphorylation and turnover.

Previous studies link PTP1B to HER2-driven tumorigenesis and survival under hypoxic stress, including RNF213-dependent regulation of non-mitochondrial oxygen consumption (40, 41, 54, 55). HSP90α stabilizes HIF-1α and oncogenic clients such as HER2 and AKT, whereas CFL1 regulates actin dynamics and participates in tumor progression and apoptotic responses (65–74, 78, 79). These functions provide a plausible context for the reduced survival signaling and increased apoptosis observed following PTP1B targeting. HSP90 inhibition also reduced viability in resistant cells, consistent with an HSP90-associated vulnerability. However, these observations do not formally establish that loss of HSP90α or CFL1 mediates the effects of PTP1B inhibition or directly connects substrate turnover to changes in HIF-1α or BCL2. Our mechanistic analyses focused on AKT and ERK, but trastuzumab resistance can also involve compensatory signaling through EGFR/HER3, IGF-1R, MET, and SRC–EphA2 (26–30, 84). These pathways may cooperate with, or function independently of, the PTP1B-dependent network identified here.

Together, our findings identify PTP1B as a therapeutic vulnerability in trastuzumab-resistant HER2-positive breast cancer models and support a working model linking substrate stability to survival signaling (Figure 8D). Developing PTP1B-directed therapeutics has historically been challenging because the conserved, charged catalytic site complicates selectivity and cellular penetration, while adequate exposure and a therapeutic window remain essential for clinical translation. Advances in allosteric inhibition, structural understanding, and medicinal chemistry, together with the progression of PTP-directed agents into clinical evaluation, mark an encouraging stage in phosphatase drug development. These developments create opportunities to translate mechanistic insights into treatments for cancers resistant to existing therapies. In trastuzumab-resistant HER2-positive breast cancer, our results provide a preclinical rationale for evaluating PTP1B inhibition as an additional therapeutic approach. Defining biomarkers of dependency, confirming target engagement, and establishing efficacy and tolerability, alone or in combination with HER2-directed treatments, will be important steps toward determining whether this strategy can benefit patients whose tumors no longer respond to trastuzumab.

## Acknowledgements

NKT is the Caryl Boies Professor of Cancer Research at Cold Spring Harbor Laboratory. We are grateful to Andreas Grill (DepYmed Inc.) for providing PTP1B inhibitors. We thank Dr. Timothy Haystead (Duke University) for generously sharing the HSP90 inhibitor, SNX5422. We thank J. Habel and staff at the Animal Shared Resource of the Cold Spring Harbor Laboratory Cancer Center for animal procedures, housing, and maintenance support. We thank James Rouse (CSHL Bioinformagics) and Hardik Patel (CSHL Organoid core) for their input. VM thanks Dr. David Tuveson, Dr. Leemor Joshua-Tor, and Dr. Sydney Gary for their advice and support.

## Author contributions

Conceptualization: NKT, VM

Data curation, analysis and investigation: VM, MCP, CF, PC

Methodology: VM, MCP, CF, SR, SK, PN

Funding acquisition and Resources: NKT

Project administration: NKT

Supervision: NKT and VM

Writing: NKT and VM, with review and editing by all authors.

All authors have approved the submitted version of the manuscript

## Materials Methods

### Antibodies and reagents

Antibodies and blotting reagents used in this study include the following: Mouse monoclonal anti-FLAG (M2, F3165 and F1804; MilliporeSigma); anti-tubulin (B-5-1-2, T5168; MilliporeSigma); anti-vinculin (hVIN-1, V9131; MilliporeSigma); anti-pTyr (4G10, 05-321; MilliporeSigma); Anti-Pyr−Agarose antibody (PT-66,A1806, MilliporeSigma) anti-pTyr conjugate agarose (4G10, 16-101; MilliporeSigma); anti-pTyr sepharose bead conjugate (pTyr-100, 9419; Cell Signaling Technologies, CST);; anti-PTP1B (FG6, Tonks Lab; CSHL) anti-HER2/ERBB2 (29D8, 2165; CST); SH2S-Magic Agarose beads (Precision proteomics) anti-phospho-p44/42 MAPK(Erk1/2)(Thr202/Tyr204) (E10, 9106S; CST); anti-p44/42 MAPK(Erk1/2) (137F5, 4695; CST); anti-CFL1 (5175, CST), p-CFL1 (PA5-105535, Thermo Scientific), anti-Hsp90-a1 (8165, CST), anti-HIF-a1 (NB100-449, Novus Biologicals), anti-Cyclin D1 (2922, CST), Rabbit polyclonal anti-phospho-Akt (Ser473) (9271; CST); anti-Akt (9272; CST); Goat polyclonal anti-mouse-IgG(H+L)-HRP conjugate (155-035-003; Jackson ImmunoResearch Inc); anti-mouse-IgG(H+L)-AlexaFluor488 conjugate (155-545-003; Jackson ImmunoResearch Inc); anti-rabbit-IgG(H+L)-HRP conjugate (111-035-003; Jackson ImmunoResearch Inc); Donkey polyclonal anti-rabbit-IgG(H+L)-AlexaFluor568 conjugate (A10042; ThermoFisher Scientific); and streptavidin-HRP conjugate (95017-426; GE Lifesciences). All antibodies were used at dilutions recommended by their manufacturers or empirically optimized. Reagents used throughout this study were purchased from MilliporeSigma unless otherwise specified. All liquid chromatography (LC) and MS solutions/solvents used were of Optima LC/MS grade from Fisher Scientific. sgRNA PTP1B#1-5′-GATGTAGTTTAATCCGACTA-3′. sgRNA PTP1B#2-5′-GGCCCTTTGC CTAACACATG-3′. sgRNA PTP1B#3-5′-GACCGCATGTGTTAGGCAAA-3′. sgRNA ERBB2 L755S – 5′-CAGTGGCCAT CAAAGTGTTG-3′. sgRNA ERBB2 V777L – 5′-AGCATACGTGATGGCTGGTG-3′. PTP1B knockout efficiency (≈90%) in SKBR3-HR cells was confirmed by ICE analysis of Sanger sequencing data. Primers used for Sanger sequencing to assess knockout efficiency are indicated: forward 5′-ggtatgctgaaatcctgaccttctaaag-3′, reverse-5′-ctaatctcaactaaaacagggcttcagc-3′, forward 5′-ggtagaaaatggagctgcagttact –3′, reverse 5′-tacgtaaaggaaatacaaaggcaag-3′, forward 5′-ctacgtaaaggaaatacaaaggcaag –3′, reverse 5′-ggtagaaaatggagctgcagttactg –3′. Primers used for Sanger sequencing to confirm the ERBB2 mutations: L755S: forward 5′-Gcagttacagcggagaaggga –3′, reverse 5′-aaatagtctaggtttgcgggagtc –3′, V777L: forward 5′-gcccggcctaatctttgtatttt –3′, reverse 5′-aatctgcatacaccagttcagc –3′

### Cell lines and culture conditions

Human embryonic kidney– and breast cancer-derived cell lines (ATCC) were cultured as a monolayer and maintained at 37 °C with 5% CO2 in Dulbecco’s Modified Eagle’s Medium (HEK293T and BT474) or Ham’s F-12 Dulbecco’s Modified Eagle’s Medium (JIMT1), supplemented, 2 mM L-glutamine, 100 units/mL penicillin, 100 μg/ml streptomycin (Gibco), and 10% (v/v) fetal bovine serum (Atlas Biologicals). HCC 1419, HCC1569 & HCC1954 cell lines obtained from ATCC were cultured in RPMI (ATCC) with 10% FBS, supplemented with penicillin-streptomycin. Cells were authenticated and tested for mycoplasma contamination (CSHL Tissue Culture Facility Shared Resource). Trastuzumab (Herceptin) –resistant cell lines were generated, similar to previously described methods (54, 60). Briefly, cells were selected for long-term outgrowth through continuous culture in Herceptin-containing medium for ∼6 to 9 months at 32 μg/ml Trastuzumab. Experiments involving resistant cells were carried out on Polyclonal trastuzumab-resistant cells generated from these outgrowth conditions as a pool of resistant cells continuously cultured in media supplemented with Herceptin, or with the de novo-resistant human breast cancer cell lines JIMT1 (87), HCC1569, HCC1419, and HCC1954. All experiments were carried out with cells grown to 70 to 80% confluence, free of mycoplasma. Trastuzumab (Herceptin) was a generous gift from Northwell Health. All cells were cultured at 37 °C with 5% CO2.

### Patient-Derived Organoid (PDO) Culturing

HER2+ tumor organoid lines were previously characterized at the genomic and transcriptomic levels (85). Organoids were cultivated in a specialized medium (REF) composed of 10% R-Spondin1 conditioned medium, 5 nmol/L Neuregulin 1 (Peprotech, 100–03), 5 ng/mL FGF7 (Peprotech, 100–19), 20 ng/mL FGF10 (Peprotech, 100–26), 5 ng/mL EGF (Peprotech, AF100–15), 100 ng/mL Noggin (Peprotech, 120–10C), 500 nmol/L A83–01 (Tocris Bioscience, 2939), 5 μmol/L Y-27632 (AbMole, Y-27632), 500 nmol/L SB202190 (Sigma-Aldrich, S7067), 1x B27 supplement (Gibco, 17504–44), 1.25 mmol/L N-acetylcysteine (Sigma-Aldrich, A9165), 5 mmol/L nicotinamide (Sigma-Aldrich, N0636), 50 μg/mL Primocin (Invitrogen, ant-pm-1), 1x GlutaMAX (Invitrogen, 12634– 034), 10 mmol/L HEPES (Invitrogen, 15630–056), and 100 U/mL penicillin/streptomycin (Invitrogen, 15140–122) in 1x advanced DMEM-F12 (Invitrogen, 12634-034). The cultures were passaged every 6 to 12 days using TrypLE (Thermo Fisher Scientific, 12605028), facilitating the breakdown of organoids into smaller cell clusters, which were subsequently replated in Matrigel. The HCM-CSHL-0155 organoid line is available from ATCC.

### Generation of PTP1B-KO Cells

PTP1B knockout cell lines (SKBR3-HR, BT474-HR, JIMT1, HCC1569 & HCC1419) were generated using CRISPR/Cas9 ribonucleoprotein (RNP) delivery via the Lonza 4D-Nucleofector System. Multiple single guide RNAs (sgRNAs) targeting an early coding exon (3) of PTP1B were designed using the Synthego Design Tool and CHOPCHOP and synthesized commercially. RNP complexes were assembled by incubating recombinant Streptococcus pyogenes Cas9 V2cut protein with sgRNA on ice for 10–15 min. Cells (10⁶ per reaction) were harvested during logarithmic growth, washed with phosphate-buffered saline, and resuspended in nucleofection solution (Lonza). The cell suspension was mixed with RNP complexes and electroporated using cell–type–specific programs according to the manufacturer’s instructions. Immediately after nucleofection, cells were transferred into pre-warmed complete growth medium and allowed to recover for 48–72 h. Individual clones were expanded and screened for gene disruption by PCR amplification of the target locus followed by Sanger sequencing. Loss of PTP1B expression was confirmed by immunoblot analysis.

### Cell proliferation and viability assays

Cell proliferation was determined by cell count on a hemocytometer. Briefly, cells were seeded into 6-well culture plates at 1.25 × 105 cells/well. The following day, the media was exchanged with media containing either vehicle, 32 μg/ml Herceptin, or 2 μM DPM-1003. At designated time points, cells were washed in PBS, trypsinized, and counted. Cell viability was determined by an ATP-dependent luminescence assay (CellTiter-Glo; Promega) as per the manufacturer’s instructions. Briefly, cells were seeded into white opaque 96-well culture plates at 1 × 104 cells/well. Media was exchanged the following day, to media containing either vehicle, Herceptin, or DPM-1003 as above. After 72 h, viability was measured by luminescence on a SpectraMax i3x multi-Mode plate reader (Molecular Devices). Data were collected from n = 3 independent experiments (biological replicates) performed individually in at least triplicate (technical replicates). Statistics were analyzed using an unpaired t-test. ns, not significant; ∗p < 0.05, ∗∗p < 0.01, ∗∗∗p < 0.001. Caspase-3/7 activity was measured using the Caspase-Glo® 3/7 Assay (Promega) according to the manufacturer’s instructions. Cells or tumor organoids were seeded in white-walled 96-well plates and treated as indicated above. Following treatment, an equal volume of Caspase-Glo® 3/7 reagent was added to each well after 72 h, and plates were incubated at room temperature for 30 min in the dark. Luminescence was measured using a SpectraMax i3x multi-Mode plate reader (Molecular Devices), and caspase-3/7 activity was compared to untreated controls in percent as indicated.

### Immunoblotting

Briefly, cells were washed with ice-cold PBS, collected and lysed in modified RIPA lysis buffer with additives (50 mM HEPES pH 7.4, 150 mM NaCl, 1% NP-40, 0.5% sodium deoxycholate, 0.1% SDS, 1 mM EDTA, 1× cOmplete protease and Phosphatase inhibitor cocktail (Roche), 1 mM PMSF, Incubate samples on ice for 10min, pipetting out u samples every 5min using and then centrifuged at 15,000xg for 15 min at 0 °C. Soluble lysates were quantified using the Bradford protein assay kit (Thermo Fisher Scientific #23200). 5X SDS loading dye was added to protein lysates and heated at 95 °C for 5 minutes. 25-50ug of whole-cell lysates were resolved by 10% or 15% SDS-PAGE, transferred to nitrocellulose membranes (Millipore Sigma), then membranes were blocked with 5% BSA (Equitech-Bio) in TBS containing 0.1% Tween-20 (TBST) for 1 h at room temperature with gentle orbital shaking. Membranes were incubated with primary antibodies in 5% BSA/TBST overnight at 4 °C, then washed with TBST, and incubated with HRP-conjugated secondary antibodies in 5% skim milk/TBST (Lab Scientific) for 1.5 h at room temperature, all with gentle platform rocking. Membranes were washed with TBST, then visualized by chemiluminescence on autoradiographic film using SuperSignal Western Blot Enhancer Substrate (ThermoFisher Scientific).

### Co-immunoprecipitation and Substrate Trap Experiments

For the Hsp90a or CFL1 degradation assay, the SKBR3-HR-Wild type (WT) and SKBR3-HR-PTP1B knockout cells were transfected with an HA-Ubiquitin plasmid from Addgene (Catalog #18712). The SKBR3-HR-WT and KO expressing HA-ubiquitin were lysed using RIPA buffer and were pulled down using Cell Signaling Anti-Ubiquitin Magnetic Beads (Cell Signaling). The cells were then washed three times using RIPA buffer using a DynaMagTM-2 Magnet rack (ThermoFisher, Catalog # 12321D). Elution from the beads was done by adding 50 μL of 2 X SDS loading dye to 50 ull of beads and heating the mixture for 10 mins at 50 ° C. The beads were then pelleted, and the supernatant was transferred to a new tube. The elution step was repeated twice, and the samples were pooled together. The samples were run on an immunoblot and probed using antibodies against Hsp90a or CFL1. Substrate trapping experiments were performed as described previously (49,53). HEK293T cells were cultured in DMEM supplemented with 10% FBS at 37°C and 5% CO2 in a humidified atmosphere. For transfection, FLAG-tagged WT PTP1B and PTP1B mutant (PTP1B-D181A) were used. Cells were transfected with Lipofectamine 2000 (Life Technologies) for 36 hours following the manufacturer’s instructions. Flag-tagged PTP1B was immunoprecipitated using anti-FLAG magnetic agarose (Thermo Fisher Scientific, #A36797). The resulting complexes were analyzed by immunoblotting.

### Mass Spectrometry Phosphoproteomics Sample Preparation

All reagents and solvents used were MS-grade. Cells were lysed in 5% SDS/100 mM triethylammonium bicarbonate (TEAB) containing 2x protease inhibitor and 1x phosphatase inhibitor via probe sonication. Cysteine residues were reduced with 3 mM tris(2-carboxyethyl) phosphine and alkylated with 10 mM iodoacetic acid. A 5mg aliquot of each sample was proteolyzed using the S-Trap method (Protifi). In short, proteins were emulsified with 1.2% phosphoric acid, diluted 6x in binding/wash buffer (90% methanol/100mM TEAB), loaded on respective S-traps, washed five times with 0.5 mL binding/wash buffer, and proteolyzed using 60 μg sequencing-grade modified porcine trypsin (Promega) overnight at 37 °C. Peptides were eluted 3x with 70% ACN/100mM TEAB. Each sample was labeled with the respective TMT-10plex reagents (Pierce) for 1 hour (RT) and the reactions were quenched with 5% hydroxylamine. Complete TMT labeling and equimolarity of the pool was checked before affinity enrichment. TMT-encoded peptide samples were combined, lyophilized, and desalted by C18 solid-phase extraction. Eluates were divided into 5 aliquots for subsequent parallel affinity enrichments: 1/2 was used for pTYR enrichment, 1/50th was aliquoted for TiO2 enrichment, 1/50th was aliquoted for FeNTA enrichment, 1/4 was aliquoted for PT66/4G10 enrichment, and the remainder was used for SH2 enrichment. FeNTA and TiO2 enrichments were conducted following the manufacturer’s protocol (Pierce). The pTYR enrichment using the PTMScan HS Phospho-Tyrosine (P-Tyr-1000, Cell Signaling Technologies, part# 38572) was conducted using the manufacturer’s protocol, except that the sample was incubated with the beads overnight at 4^0^C with end-over-end mixing. The SH2 IP was also conducted according to the manufacturer’s instructions. For the PT66/4G10 enrichment, the sample was resuspended in 50mM Tris buffer/100mM NaCl/2x protease inhibitors/1x phosphatase inhibitors. 4G10-conjugated beads were mixed with PT66 beads before washing the beads with cold PBS and cold Tris buffer + inhibitors. The sample was mixed with the PT66/4G10 beads and allowed to incubate overnight at 4C on a rotator. The flow-through was moved to a new tube, and the beads were washed with cold Tris buffer + inhibitors, then cold water, before eluting with 0.2% TFA. 1/10th of each eluate from phosphopeptide enrichment was then analyzed by LC/MS/MS.

### LC/MS analysis

Peptides were loaded via PepMapTM Neo Trap Cartridge (Thermo Part #: 174500) onto a 30 cm x 75 μm ID analytical column packed in-house with Reprosil 1.9 μm C18 silica particles (Dr. Maischt), and resolved on a 5-35% acetonitrile gradient in water (0.1% formic acid) at 200 nl/min over 120 min. Eluting peptides were ionized by electrospray (2200V) using a 10 μm ID silica emitter (FossilIon) and transferred into a Thermo Scientific Exploris480 quadrupole-orbitrap (data-dependent analyses) or Fusion Lumos (data-dependent and targeted) tribrid mass spectrometer (MS). For data-dependent analyses, the MS was set to collect 120,000-resolution precursor scans (m/z 380-2000 Th, automatic gain control set to 500,000 charges, max accumulation time=50 ms) every 3 s. Precursor ions were selected for fragmentation in data-dependent mode with a dynamic exclusion list (10 ppm mass tolerance, 60 s exclusion). HCD fragmentation was performed at stepped 28,33,38% (Exploris480) or 30,35,40% (Fusion Lumos) normalized energy, with spectra collected in the Orbitrap at 60,000 resolutions (first mass locked to 100 Th. 50 ms max accumulation time, 50,000 charges max) in centroid mode.

### MS Data analysis

Spectra were matched against the UniProt human database (as of April 2023) and a database of common contaminants (cRAP), using the Mascot scoring function (ProteinMetrics) within ProteomeDiscoverer (Thermo). K-TMTpro, Nterm-TMTpro, STY-phosphorylation, M-oxidation and N/Q-deamidation were set as variable modifications. Carboxymethylation was set as a fixed adduct on Cys residues. Mass tolerance set at 5 ppm for MS1, and 0.02 Da for MS2. Peptide-spectral matches and peptide and protein identifications were filtered to maintain FDR<1% using the target-decoy approach within Percolator. TMT reporter ion intensities recorded for each identified peptide were used for relative quantification across samples and used as a quantitative metric for protein relative quantification. Peptide-level identifications were used to evaluate differential phosphorylation across the detected sites. In total, 26,045 peptides were identified at FDR <0.01 in the enriched samples. After filtering out peptides with no quantitative value, 20,867 peptides were used for quantitative analysis, of which 6,084 peptides were phosphorylated. 786 of these were identified as phosphorylated at tyrosine residues, or the phosphorylation could not be localized based on MS2, but the peptide sequence contained a tyrosine residue. Peptide TMT abundances were normalized using the normalization factors from the protein-level screen results. To evaluate differential abundance of phosphorylated peptides for each experimental condition, the ratio of the mean reporter ion intensities in different samples was compared, and the statistical significance of the difference between the means was assessed using Student’s t-test. Volcano plots for the different experimental conditions are shown below. P-values were calculated in Perseus using a two-sample Student’s t-test and a threshold p-value of 0.05.

### Mammary fat pad xenografts and drug treatments

All animal procedures were performed in accordance with Institutional Animal Care and Use Committee (IACUC) guidelines and were approved by the institutional ethics committee. Female immunodeficient mice (4–6 weeks old; athymic nude or NOD/SCID) were obtained from The Jackson Laboratory and maintained under specific pathogen-free conditions. Orthotopic mammary fat pad xenografts were established using SKBR3-HS, SKBR3-HR, or JIMT1 cells. Cells were harvested during logarithmic growth, washed twice with sterile PBS, and resuspended in a 1:1 mixture of PBS and Matrigel (Corning). A total of 1 × 10⁶ cells in 100 µL were injected into the fourth inguinal mammary gland fat pad as previously described (86).

For JIMT1 xenografts, drug treatment was initiated one week after cell implantation. Mice received vehicle control, trastuzumab (Herceptin; 15 mg/kg, intraperitoneally), or DPM-1003 (35 mg/kg orally or 10 mg/kg intraperitoneally), administered twice weekly. Tumor growth and animal health were monitored regularly, and tumor dimensions were measured twice weekly using digital calipers. Tumor volume was calculated as (length × width²)/2. Animals were euthanized when tumors reached 2 cm³ or developed ulceration, in accordance with institutional guidelines. At the indicated experimental endpoints, tumors were excised, weighed, and snap-frozen in liquid nitrogen for downstream molecular analyses. Data were obtained from 11 animals per treatment group across three independent experiments.

### Statistical analysis

All data are presented as mean ± SD and represent a minimum of three independent biological replicates unless otherwise indicated. For endpoint analyses, including CellTiter-Glo (CTG) and Caspase-Glo 3/7 apoptosis assays and immunoblot quantification, statistical significance was determined by one-way or two-way ANOVA, as appropriate, followed by Bonferroni’s multiple-comparisons test. For longitudinal analyses, including cell proliferation and tumor growth curves, statistical significance was determined by two-way ANOVA or mixed-effects analysis, as appropriate, followed by Tukey’s multiple-comparisons test. The specific statistical test used for each experiment is indicated in the corresponding figure legend. Statistical significance is denoted as follows: ns, not significant; */#P < 0.05; **/##P < 0.01; and ***/###P < 0.001.

## Supplementary Figure Legends

**Supplementary Figure S1.**
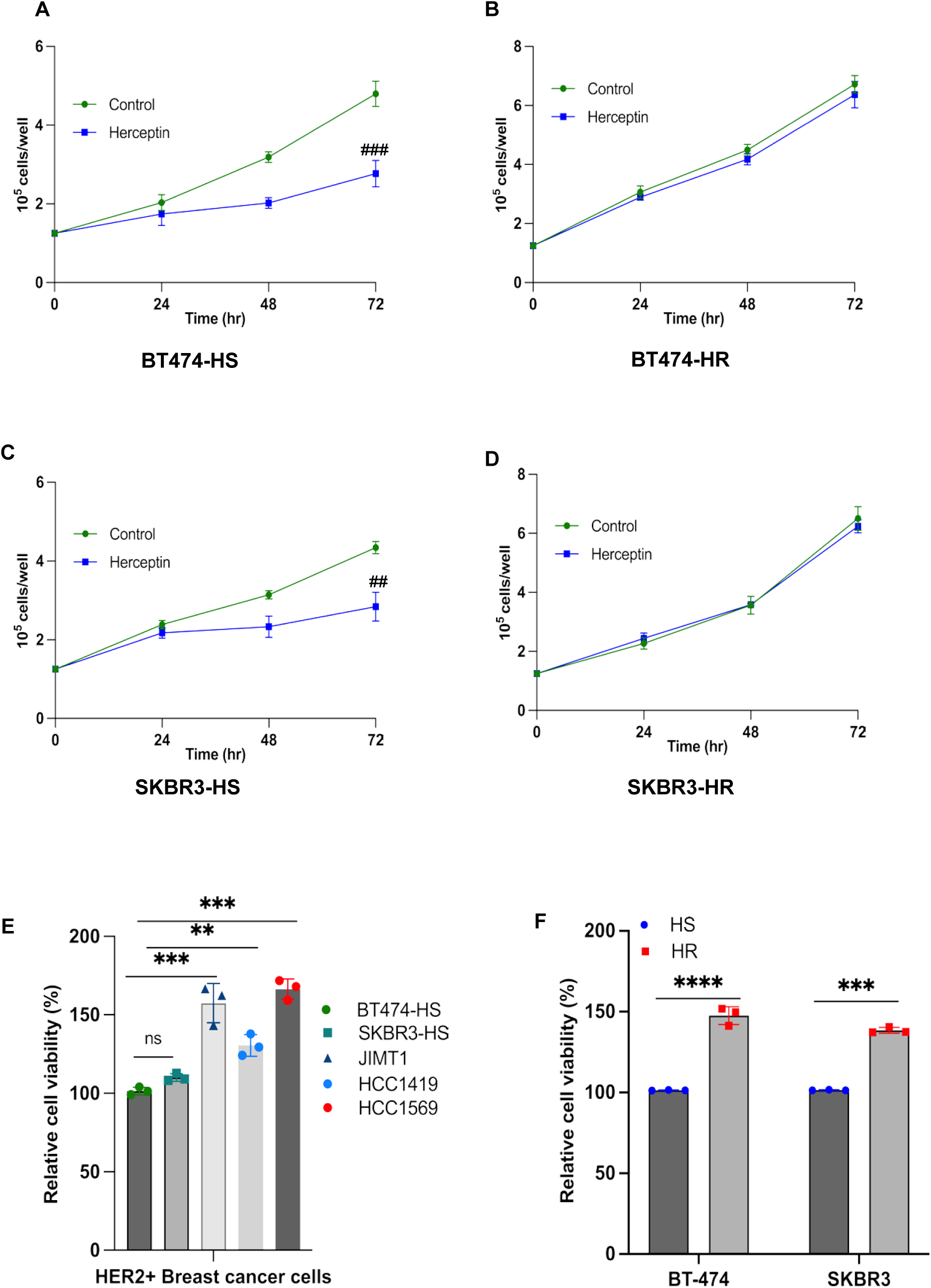
Generation and validation of acquired trastuzumab-resistant HER2-positive breast cancer cell models. (**A–D**) Cell proliferation was assessed by direct cell counting using a hemocytometer in trastuzumab-sensitive BT474-HS (A) and SKBR3-HS (C) cells and their corresponding acquired trastuzumab-resistant derivatives, BT474-HR (B) and SKBR3-HR (D), following treatment with vehicle or trastuzumab (32 μg/mL) for 72 h. Trastuzumab significantly suppressed proliferation in the sensitive parental cells but had little effect on their resistant derivatives, confirming the acquisition of trastuzumab resistance. **(E)** Cell viability measured by the CellTiter-Glo (CTG) assay in trastuzumab-sensitive (HS; BT474 and SKBR3) and intrinsic trastuzumab-resistant (HR; JIMT1, HCC1419, HCC1569, and HCC1954) HER2-positive breast cancer cell lines, demonstrating increased proliferative capacity in resistance cells. **(F)** CTG assay comparing parental trastuzumab-sensitive (BT474-HS and SKBR3-HS) cells with their corresponding acquired resistance (BT474-HR and SKBR3-HR) cell lines, demonstrating enhanced proliferation of resistance cells. Data are presented as mean ± SD from three independent biological experiments unless otherwise indicated. Data were normalized to the respective control for visualization. Statistical significance was determined using one-way ANOVA (E) or two-way ANOVA (A, C and F) followed by Bonferroni’s multiple comparisons test. ns, not significant; */ # P < 0.05; **/## P < 0.01; ***/###P < 0.001.

**Supplementary Figure S2.**
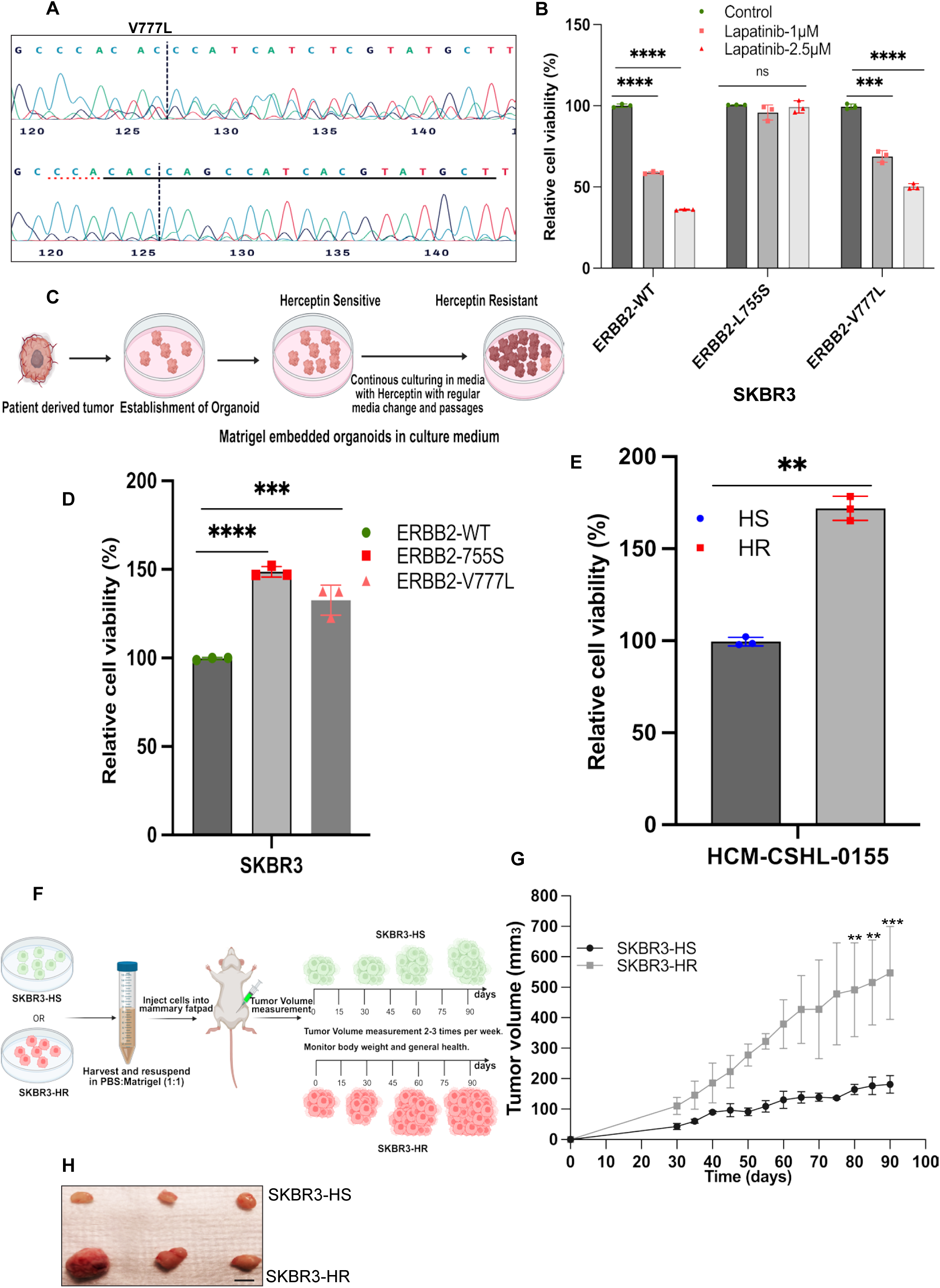
Characterization of ERBB2-mutant, trastuzumab-resistant cell, organoid, and xenograft models. **(A)** Sanger sequencing chromatograms confirming successful CRISPR–Cas9-mediated introduction of the endogenous ERBB2 V777L mutation into SKBR3 cells. **(B)** Cell viability measured by CellTiter-Glo (CTG) assay in SKBR3 cells expressing wild-type (WT) ERBB2 or the endogenous L755S and V777L mutant variants following lapatinib treatment (72 h). Cells expressing the L755S mutation exhibited no sensitivity to lapatinib compared with WT and V777L cells. Data are presented as mean ± SD from three independent biological experiments unless otherwise indicated. Data were normalized to the respective control for visualization. Statistical significance in Figure B was determined using two-way ANOVA followed by Bonferroni’s multiple comparisons test. ns, not significant; *P < 0.05; **P < 0.01; ***P < 0.001; ****P < 0.0001. **(C)** Experimental scheme illustrating the generation of HER2-positive trastuzumab-resistant (HR) tumor organoids from parental trastuzumab-sensitive (HS) organoids by prolonged trastuzumab exposure. Created in BioRender. M, V. (2026). https://BioRender.com/sw1qf8n **(D)** CTG assay comparing SKBR3 cells expressing wild-type (WT) **ERBB2** or the endogenous L755S and V777L mutant variants, demonstrating increased proliferation of mutant cells relative to WT controls. **(E)** CTG assay comparing the growth of HER2-positive tumor organoids derived from trastuzumab-sensitive (HCM-CSHL-0155-HS) and trastuzumab-resistant (HCM-CSHL-0155-HR) lines. **(F)** Experimental scheme illustrating orthotopic xenograft studies using SKBR3-HS and SKBR3-HR cells. Mice were implanted orthotopically into the mammary fat pad with **5 × 10⁶** cells, and tumors were harvested when they reached approximately **600–700 mm³**. Created in BioRender. M, V. (2026) https://BioRender.com/sw1qf8n. **(G)** Tumor growth curves of SKBR3-HS and SKBR3-HR xenografts showing increased tumor growth in the trastuzumab-resistant group. (H) Representative images of orthotopic xenograft tumors derived from SKBR3-HS and SKBR3-HR cells, illustrating enhanced tumor growth in the resistant group. Scale bar, 1 cm. Data are presented as mean ± SD from three independent biological experiments unless otherwise indicated. Data were normalized to the respective control for visualization. Statistical significance was determined using one-way ANOVA (C and D) or two-way ANOVA (B and F), followed by Bonferroni’s multiple comparisons test. ns, not significant; *P < 0.05; **P < 0.01; ***P < 0.001; ****P < 0.0001.

**Supplementary Figure S3.**
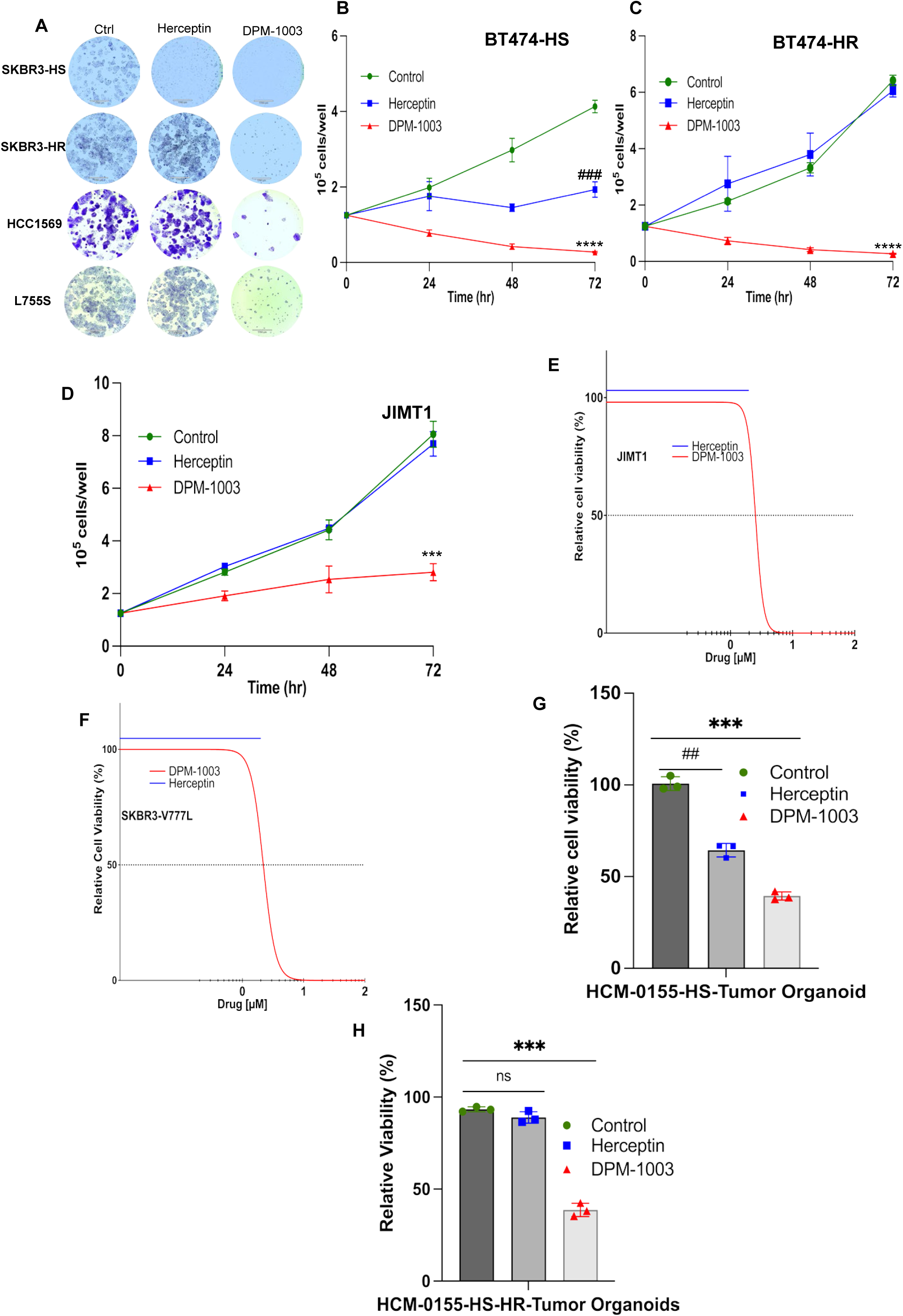
Pharmacological inhibition of PTP1B suppressed growth in trastuzumab-resistant HER2-positive breast cancer models. **(A)** Representative colony formation assay images demonstrate that PTP1B inhibition markedly reduced growth in all cell models, whereas trastuzumab showed inhibition only in sensitive cells. **(B–D)** Cell proliferation measured by hemocytometer counting in BT474-HS (B), BT474-HR (C), and JIMT1 (D) cells following treatment with vehicle, trastuzumab (32 μg/mL), or the PTP1B inhibitor DPM-1003 (2 μM) for 72 h. DPM-1003 suppressed proliferation in trastuzumab-resistant cells, whereas trastuzumab showed no effect. **(E–F)** Cell viability was measured by the CellTiter-Glo (CTG) assay following 72 h of treatment with increasing concentrations of DPM-1003 or trastuzumab in JIMT1 (E) and SKBR3 cells expressing the endogenous ERBB2 V777L mutation (F). DPM-1003 markedly reduced cell viability in both resistant models, whereas trastuzumab showed no activity. **(G–H)** Cell viability was measured by CTG assay in HER2-positive trastuzumab-sensitive (HCM-CSHL-0155-HS) (G) and trastuzumab-resistant (HCM-CSHL-0155-HR) (H) tumor organoids following treatment with trastuzumab or DPM-1003. DPM-1003 markedly reduced organoid viability in both conditions, whereas trastuzumab was effective only in the trastuzumab-sensitive organoids. Data are presented as mean ± SD from three independent biological experiments unless otherwise indicated. Data were normalized to the respective control for visualization. Statistical significance was determined by One-way ANOVA (H and I), followed by Bonferroni’s multiple-comparisons test, or by mixed-effects analysis (B–D), followed by Tukey’s multiple-comparisons test. Asterisks (*, **, ***) indicate comparisons between control and DPM-1003-treated groups; number signs (#, ##, ###) indicate comparisons between control and Herceptin-treated groups. */#P < 0.05; **/##P < 0.01; ***/###P < 0.001.

**Supplementary Figure S4.**
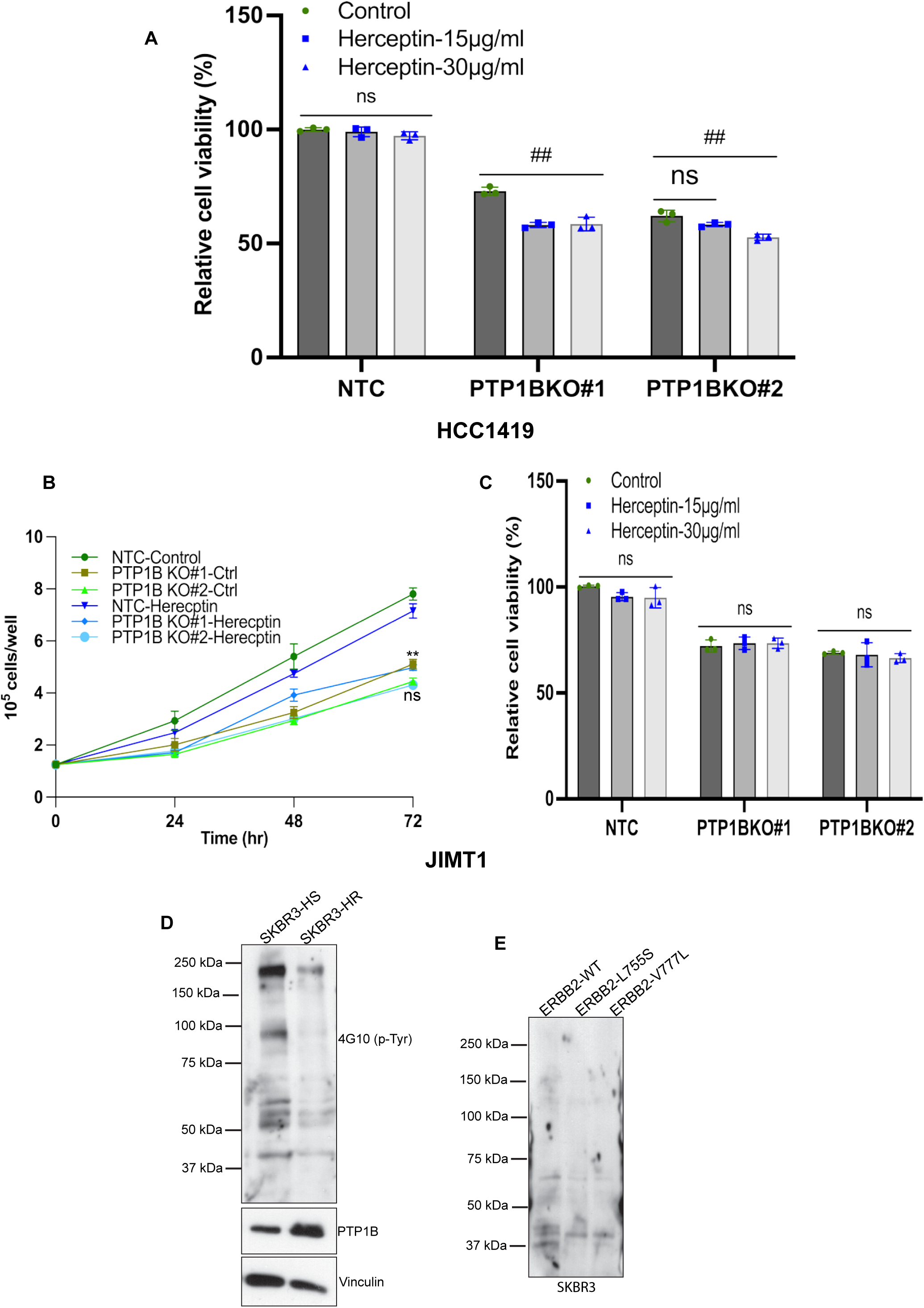
Validation of PTP1B-dependent regulation of trastuzumab sensitivity and phosphotyrosine signaling. **(A)** Cell viability measured by CellTiter-Glo (CTG) assay in HCC1419 cells expressing a non-targeting control (NTC) or *PTPN1* knockout following trastuzumab treatment (72 h), demonstrating restoration of trastuzumab sensitivity following *PTPN1* loss. **(B–C)** Cell proliferation measured by hemocytometer counting of JIMT1-PTP1B-KO cells (B) and CTG cell viability (C) assays of the same cells expressing a non-targeting control (NTC) or *PTPN1* knockout following trastuzumab treatment (72 h), showing that *PTPN1* loss did not restore trastuzumab sensitivity in these cells. **(D–E)** Immunoblot analysis of phosphotyrosine (p-Tyr) levels in SKBR3-HR cells with elevated PTP1B (D) and SKBR3 cells expressing endogenous ERBB2 mutant variants (E), suggesting a relation between PTP1B expression and phosphotyrosine levels. Data are presented as mean ± SD from three independent biological experiments unless otherwise indicated. Data were normalized to the respective control for visualization. Statistical significance was determined by two-way ANOVA (A and C), followed by Bonferroni’s multiple-comparisons test, or by mixed-effects analysis (B), followed by Tukey’s multiple-comparisons test. Asterisks (*, **, ***) indicate comparisons between control and PTP1B-KO groups; number signs (#, ##, ###) indicate comparisons between untreated PTP1B-KO and PTP1B-KO + Herceptin-treated groups. */#P < 0.05; **/##P < 0.01; ***/###P < 0.001.

**Supplementary Figure S5:**
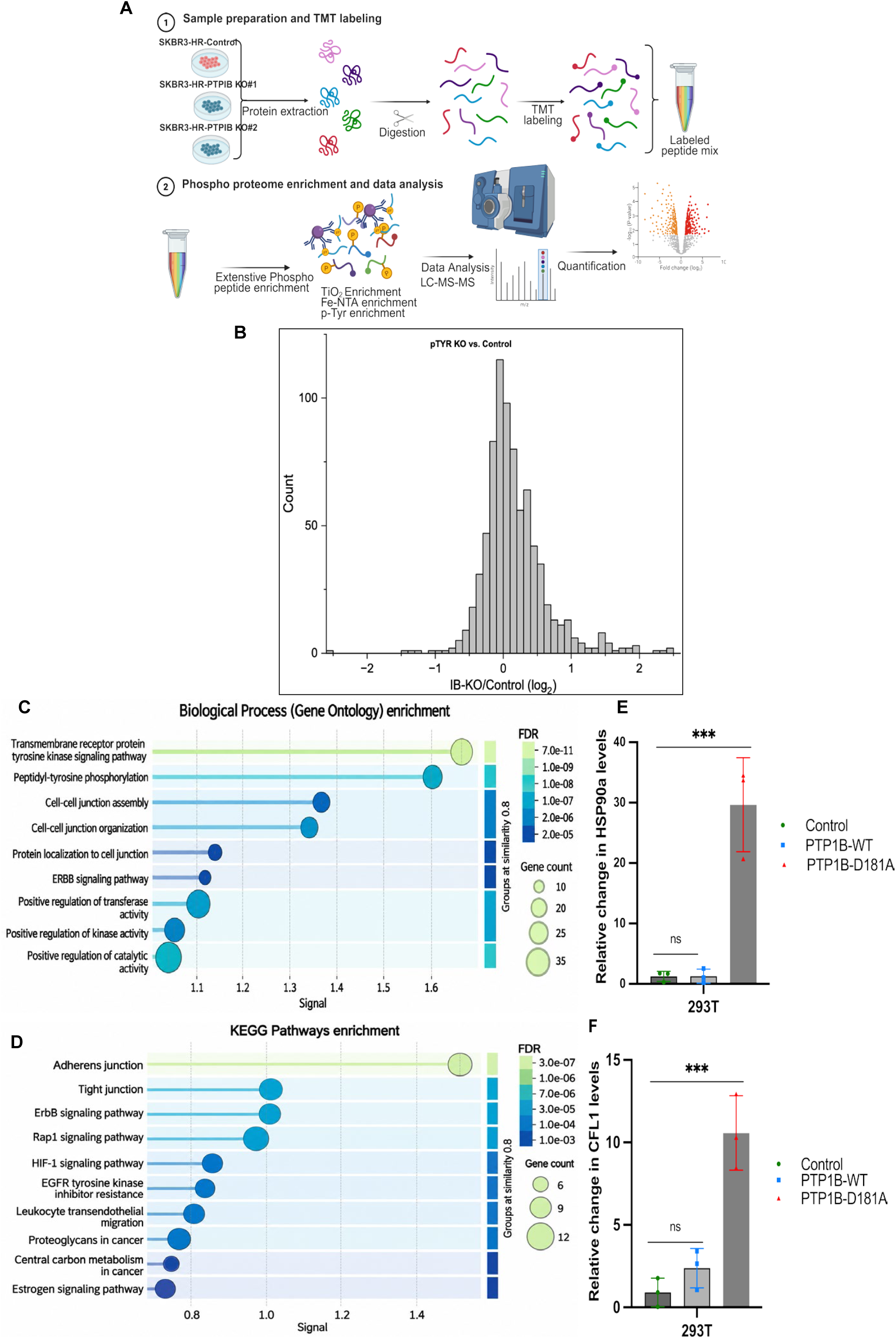
PTP1B phosphotyrosine proteomic profiling, pathway enrichment, and substrate-trapping validation. **(A)** Experimental workflow for TMT-based quantitative phosphoproteomic profiling of SKBR3-HR cells with or without PTP1B, including phosphopeptide and phosphotyrosine enrichment followed by LC–MS/MS. Created in BioRender. M, V. (2026) https://BioRender.com/sw1qf8n. **(B)** Distribution of phosphotyrosine-containing peptides identified by LC–MS/MS, showing an overall increase in phosphotyrosine abundance following PTP1B loss. **(C)** Gene Ontology (GO) Biological Process enrichment analysis of differentially regulated phosphotyrosine proteins following PTP1B loss, showing enrichment of biological processes associated with protein tyrosine phosphorylation and receptor tyrosine kinase signaling. **(D)** KEGG pathway enrichment analysis of differentially regulated phosphotyrosine proteins, identifying HER2/ERBB2 signaling, HIF signaling, and tyrosine kinase inhibitor resistance among the most significantly enriched pathways. **(E–F)** Quantification of CFL1 (E) and HSP90α (F) immunoblot signal intensities from the PTP1B substrate-trapping assay shown in Figure 5J. Data are presented as mean ± SD from three independent biological experiments unless otherwise indicated. Data were normalized to the respective control for visualization. Statistical significance was determined by One-way ANOVA (E and F), followed by Bonferroni’s multiple-comparisons test. ns, not significant; *P < 0.05; **P < 0.01; ***P < 0.001; ****P < 0.0001.

**Supplementary Figure S7.**
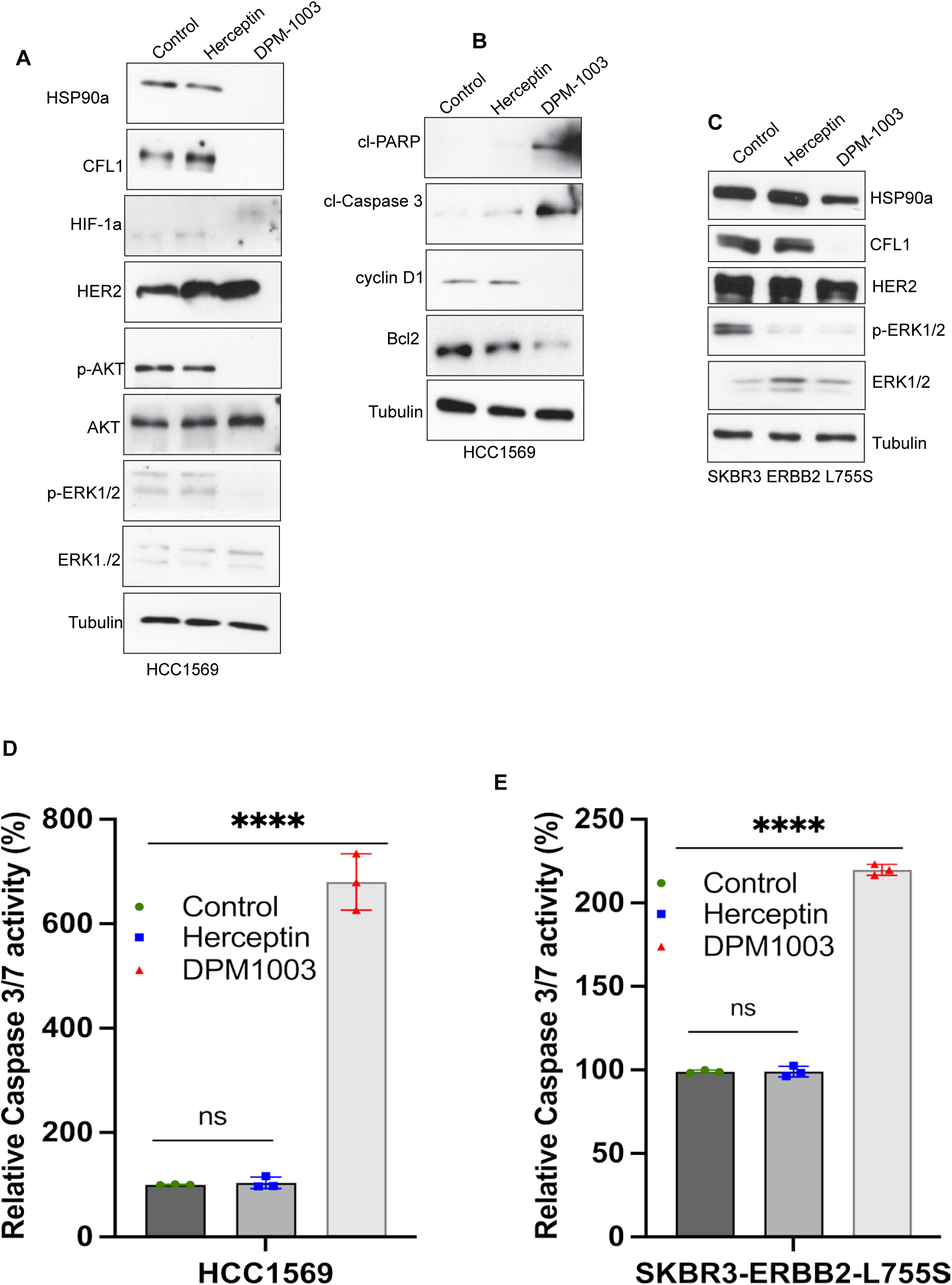
PTP1B inhibition suppressed oncogenic signaling and induced apoptosis in trastuzumab-resistant breast cancer models. **(A–B)** Immunoblot analysis of HCC1569 cells treated with vehicle, trastuzumab (32 μg/mL), or the PTP1B inhibitor DPM-1003 (2 μM) for 48 h. DPM-1003, but not trastuzumab, reduced CFL1, HSP90α1, HIF-1α, phospho-AKT, phospho-ERK1/2, BCL2, and Cyclin D1 levels while increasing cleaved PARP and cleaved caspase-3, consistent with induction of apoptosis. **(C)** Immunoblot analysis of SKBR3 cells expressing the endogenous ERBB2 L755S mutation treated with vehicle, trastuzumab (32 μg/mL), or DPM-1003 (2 μM) for 48 h. DPM-1003, but not trastuzumab, reduced CFL1, HSP90α1, and phospho-ERK1/2 levels. **(D–E)** Caspase-3/7 activity assays in HCC1569 cells (D) and SKBR3 cells expressing the endogenous ERBB2 L755S mutation (E) following treatment with vehicle, trastuzumab, or DPM-1003, demonstrating increased apoptotic activity upon PTP1B inhibition. Data are presented as mean ± SD from three independent biological experiments unless otherwise indicated. Data were normalized to the respective control for visualization. Statistical significance in D and E was determined using one-way ANOVA followed by Bonferroni’s multiple comparisons test. ns, not significant; *P < 0.05; **P < 0.01; ***P < 0.001, ****P < 0.0001.

